# Connexin 43 Regulates Intercellular Mitochondrial Transfer from Human Mesenchymal Stromal Cells to Chondrocytes

**DOI:** 10.1101/2024.03.18.585552

**Authors:** Rebecca M. Irwin, Matthew A. Thomas, Megan J. Fahey, María D. Mayán, James W. Smyth, Michelle L. Delco

**Affiliations:** Department of Clinical Sciences, College of Veterinary Medicine, Cornell University, Ithaca, NY, 14853, USA; CellCOM Research Group, Instituto de Investigación Biomédica de A Coruña (INIBIC), Servizo Galego de Saúde (SERGAS), Universidade da Coruña (UDC), A Coruña, Spain; Fralin Biomedical Research Institute at Virginia Tech Carilion, Roanoke, VA 24016, USA; Center for Vascular and Heart Research, FBRI at VTC, Roanoke, VA 24016, USA; Virginia Tech Carilion School of Medicine, Roanoke, VA 24016, USA; Department of Biological Sciences, Virginia Tech, Blacksburg, VA 24061, USA

**Keywords:** Cx43, GJA1, GJA1-20k, gap junctions, osteoarthritis, arthritis, regenerative medicine, MSCs

## Abstract

**Background:** The phenomenon of intercellular mitochondrial transfer from mesenchymal stromal cells (MSCs) has shown promise for improving tissue healing after injury and has potential for treating degenerative diseases like osteoarthritis (OA). Recently MSC to chondrocyte mitochondrial transfer has been documented, but the mechanism of transfer is unknown. Full-length connexin43 (Cx43, encoded by *GJA1*) and the truncated internally translated isoform GJA1-20k have been implicated in mitochondrial transfer between highly oxidative cells, but have not been explored in orthopaedic tissues. Here, our goal was to investigate the role of Cx43 in MSC to chondrocyte mitochondrial transfer. In this study, we tested the hypotheses that (a) mitochondrial transfer from MSCs to chondrocytes is increased when chondrocytes are under oxidative stress and (b) MSC Cx43 expression mediates mitochondrial transfer to chondrocytes.

**Methods:** Oxidative stress was induced in immortalized human chondrocytes using tert-Butyl hydroperoxide (t-BHP) and cells were evaluated for mitochondrial membrane depolarization and reactive oxygen species (ROS) production. Human bone-marrow derived MSCs were transduced for mitochondrial fluorescence using lentiviral vectors. MSC Cx43 expression was knocked down using siRNA or overexpressed (GJA1+ and GJA1-20k+) using lentiviral transduction. Chondrocytes and MSCs were co-cultured for 24 hrs in direct contact or separated using transwells. Mitochondrial transfer was quantified using flow cytometry. Co-cultures were fixed and stained for actin and Cx43 to visualize cell-cell interactions during transfer.

**Results:** Mitochondrial transfer was significantly higher in t-BHP-stressed chondrocytes. Contact co-cultures had significantly higher mitochondrial transfer compared to transwell co-cultures. Confocal images showed direct cell contacts between MSCs and chondrocytes where Cx43 staining was enriched at the terminal ends of actin cellular extensions containing mitochondria in MSCs. MSC Cx43 expression was associated with the magnitude of mitochondrial transfer to chondrocytes; knocking down Cx43 significantly decreased transfer while Cx43 overexpression significantly increased transfer. Interestingly, GJA1-20k expression was highly correlated with incidence of mitochondrial transfer from MSCs to chondrocytes.

**Conclusions:** Overexpression of GJA1-20k in MSCs increases mitochondrial transfer to chondrocytes, highlighting GJA1-20k as a potential target for promoting mitochondrial transfer from MSCs as a regenerative therapy for cartilage tissue repair in OA.

## Background

Osteoarthritis (OA) is a degenerative joint disease and the leading cause of disability in older adults, affecting >300 million people globally each year (1). OA is characterized by the degradation of articular cartilage, a highly specialized soft tissue in the joint that provides load support and a highly lubricated surface for joint articulation. Cartilage is composed of chondrocytes embedded within a dense extracellular matrix (ECM) and has little capacity for self-repair, in part due to the avascularity of the tissue. Despite decades of research, no available therapies prevent OA progression after cartilage injury (2,3), and therefore, strategies to promote cartilage health and healing hold promise toward the pressing clinical need for effective OA treatments.

Mesenchymal stromal cell (MSC)-based therapies, including extracellular vesicles (EVs) derived from MSCs (EVs-MSCs), have recently emerged as a potential regenerative therapy for OA. Injections of MSCs, or EVs-MSCs, into joints after a traumatic injury have been reported to attenuate cartilage degradation and joint degeneration in pre-clinical models (4–6). Additionally, there has been significant clinical investigation of stem cell therapies for OA with over 70 clinical trials completed and more than 40 currently underway (7). Of note, MSC injections into human OA knee joints resulted in decreased pain and cartilage catabolic biomarkers 1 year after injection (8,9). While a growing body of evidence has identified therapeutic effects of MSCs including recruitment of endogenous stem cells and immunomodulation(10–12), the exact mechanism(s) underlying the beneficial effects of MSCs on cartilage repair remains unclear. One potential mechanism is the donation of whole-organelle mitochondria from MSCs to chondrocytes. This process of intercellular mitochondrial transfer has been shown to increase when cells are stressed, and transfer can rescue injured/stressed cells by restoring cellular bioenergetics, preserving cell viability, reducing oxidative stress, and improving tissue healing across multiple cell types including the lung, heart, and brain (13–23). Recently, in orthopaedic tissues, mitochondrial transfer has been shown to improve tendon healing *in vivo* (24) and to occur from MSCs to chondrocytes (25–27).

Mitochondrial dysfunction is one of the earliest cellular responses to traumatic injury in cartilage tissue (28), and mitochondrial-targeted therapies have effectively decreased mitochondrial dysfunction and cartilage degeneration while preserving chondrocyte viability after injury *in vitro* (29,30). While these studies point to the therapeutic potential of mitochondrial transfer for cartilage repair, the mechanisms of mitochondrial transfer from MSCs to chondrocytes remain unknown. Mitochondrial transfer has been shown to occur through direct cell-cell contacts (i.e. tunneling nanotubules (TNTs)) or through microvesicles (including EVs) containing mitochondria from donor cells (21). Notably, the gap junction protein connexin43 (Cx43) has been identified as a critical regulator of both EV-and TNT-mediated mitochondrial transfer in multiple cell types (13,31–35). In the context of cartilage, both pharmacologic-(carbenoxolone disodium) and Cx43-mimetic peptide-(Gap 27) mediated inhibition of gap junctions were found to decrease mitochondrial transfer between MSCs and murine chondrocytes (25).

Cx43 is a transmembrane protein (gene name *GJA1*) that forms pores at the cell membrane (hemichannels) which can communicate with the extracellular space and/or dock with channels on opposing cells to form gap junctions effecting direct intercellular communication. When hemichannels on adjacent cells align, gap junctions are formed for direct cell-cell communication. Recently, *GJA1* was identified to undergo alternative translation to create multiple N-terminally truncated isoforms (36). Of the truncated isoforms, GJA1-20k has been implicated in mitochondrial transfer as it aids in mitochondrial motility by mobilizing mitochondria along microtubules (37), recruits actin to organize cell trafficking pathways (38), and overexpression of GJA1-20k increased mitochondrial transfer from astrocytes to neurons *in vitro* (39). Additionally, GJA1-20k is critical for trafficking Cx43 hemichannels from the Golgi apparatus to the cell membrane and could therefore support the Cx43-channel role of mitochondrial transfer as discussed above (13,31,36,40). These studies highlight the potential multi-faceted role of GJA1-20k in mitochondrial transfer, but GJA1-20k has not been investigated in mitochondrial transfer involving MSCs or chondrocytes.

The objectives of this study were to investigate the role of oxidative stress on intercellular mitochondrial transfer from MSCs to chondrocytes, and to identify the role of Cx43 and GJA1-20k in the mechanism of transfer. We hypothesize that (a) MSC to chondrocyte mitochondrial transfer will be increased when chondrocytes are under oxidative stress, (b) mitochondrial transfer from MSCs to chondrocytes will occur predominantly through direct cell contacts, (c) the magnitude of mitochondrial transfer will be dependent on Cx43 in MSCs, where MSC Cx43 knockdown will decrease mitochondrial transfer and overexpression will increase transfer, and (d) expression of GJA1-20k will strongly correlate with the rate of mitochondrial transfer.

## Methods

### Ethics Statement

The immortalized human chondrocyte cell line (T/C-28a2) was kindly provided by Dr. Miguel Otero from the Hospital for Special Surgery, New York, NY. Human MSCs were purchased from RoosterBio.

### Human Cell Culture

Human bone marrow-derived MSCs were purchased from RoosterBio (MSC-003) at passage 2. MSCs were cultured in RoosterNourish-MSC media (RoosterBio, KT-001) and incubated at 37°C and 5% CO_2_. MSCs of passage 4-6 were used in experiments. To study the role of Cx43 in mitochondrial transfer from MSCs to chondrocytes, an immortalized human chondrocyte line was used as a demonstrated model for the study of Cx43 and gap junctions (41). Chondrocytes were cultured in Dulbecco’s modified Eagle’s medium (DMEM, no glucose, L-Glutamine, and sodium pyruvate; VWR, 11966025) with 10% fetal bovine serum (R&D Systems, S11550), 1% Penicillin/Streptomycin (VWR, 100X, 97063-708), 1% sodium pyruvate (Thermo Fischer, 200 mM, 11360070), 2% L-Glutamine (Thermo Fisher, 200 mM, 25030081), 45 mg/100 mL of D-Glucose (VWR, BDH9230). Chondrocytes of passage 4-8 were used in experiments.

### Mitochondrial Membrane Polarization

JC-10 was used to quantify mitochondrial membrane potential as this stain exhibits mitochondrial potential-dependent accumulation in the mitochondria as previously described (42). Chondrocytes were seeded onto a black, flat bottomed 96-well plate at 20,000 cells per well and allowed to culture overnight. Cells were stimulated with t-BHP (0, 1, 12, 30, or 60 μM) in 1X OptiMEM for 24 hrs. FCCP is a potent mitochondrial oxidative phosphorylation uncoupler and was used as a positive control (20 μM, 20 min incubation at room temperature, Sigma Aldrich, C2920). All cells were rinsed with PBS followed by incubation with JC-10 (10 μM, Enzo Life Sciences, ENZ-52305) for 45 minutes at room temperature in the dark. Following incubation, cells were read on a plate reader with an excitation of 490 nm and 540 nm. The ratio of emission at 525/590 was calculated for each well as the ratio of depolarized/polarized mitochondria within the cells.

### Measurement of ROS

Chondrocyte ROS production was measured using the CellROX Green Flow Cytometry Assay Kit according to the manufacturer’s instructions (Invitrogen, C10492). After 24 hrs of +/-t-BHP stimulation, chondrocytes were rinsed with PBS, lifted, and incubated with CellROX Green (500 nM) for 30 minutes at room temperature in the dark. After incubation, cells were immediately analyzed using a Thermo Fisher Attune NxT flow cytometer. Unstained chondrocytes were used to set up gates.

### GJA1 Knockdown Cell Lines

siRNA was used to knockdown *GJA1* expression in human MSCs based on previous work in other cell types (43). *GJA1* siRNA (Thermo Fisher Scientific, ID HSS178257) was used to knockdown *GJA1* expression and Stealth TM RNAi (Thermo Fisher Scientific, ID 12935112) was used as a negative control. Human MSCs were plated onto 6-well plates and cultures until 60% confluent. Cells were then incubated in 1X OptiMEM (Thermo Fisher Scientific, ID 31985070) containing 100 pM of either *GJA1* siRNA or Stealth RNAi and 2 µL of Lipofectamine (Invitrogen, STEM00001) per well for 24 hrs. After incubation, media was replaced with RoosterNourish and cells were cultured for 4 days until cells were lysed for confirmation of knockdown using western blotting or used in co-culture experiments as described below.

### Cx43 and GJA1-20k Overexpression Cell Lines

Lentiviral transduction was performed to create MSCs that overexpress *GJA1* and GJA1-20k as previously described (40). Briefly, lentivirus was created from pLenti6.3-*LacZ* (control), pLenti6.3-h*GJA1*, pLenti6.3-GJA1-20k according to manufacturer instructions (Thermo Scientific, ViraPower Lentiviral Expression System). Viruses were titered and used to infect MSCs on 6-well plates in RoosterGEM (RoosterBio, M40200). After 24 hrs, fresh RoosterNourish media replaced the lentivirus media and cells were cultured for an additional 24 hrs. Cells were selected using 10 μg/ml blasticidin added to the medium. After a control well of non-transfected cells were killed by the blasticidin treatment, cells were expanded and screened for overexpression by western blotting and immunofluorescence.

### Fluorescent Cell Labeling

MSCs and chondrocytes were fluorescently labeled to distinguish between cell types during imaging and to identify mitochondrial transfer events. Lentiviral transduction was used to fluorescently label each cell type instead of using exogenous stains that could be pumped out and transferred between cells. MSCs and chondrocytes were fluorescently labeled using mitochondrial or cytoplasm targeted lentiviruses driven by an EF1α promoter for co-cultures (Takara Bio USA, Inc.). Mitochondria were labeled with GFP or mCherry (0017VCT, 0024VCT) and chondrocyte cytoplasm was labeled with mCherry (0037VCT). Chondrocytes and MSCs were transduced for 24 hrs in RoosterGEM as described above.

### Western Blotting

Cells were lysed in RIPA buffer (Pierce, 89900) supplemented with the HALT Protease and Phosphatase Inhibitor Cocktail (Thermo Scientific, 87786). Western blotting was performed as previously described (40). Briefly, cells were scraped into the RIPA buffer + HALT cocktail and centrifuged at 10,000 x g for 20 minutes at 4°C. Supernatant was collected and stored at −80°C. Protein concentration was quantified using the Bio-Rad DC Protein Assays with 10-15 μg of protein loaded for each sample. NuPage 4X Sample Buffer (Thermo Scientific) supplemented with dithiothreitol (DTT, 400 mM) was added and samples were heated for 10 min at 70°C before SDS-PAGE. Protein was transferred to a PVDF membrane using the iBlot 2 system (Thermo Fisher, IB21001). Western blotting was performed with rabbit anti-Cx43 (1:5000; Sigma, C6219) and mouse anti–α-tubulin (1:5000; Abcam, ab7291) as the primary antibodies. The anti-Cx43 antibody targets the C-terminal domain and will therefore identify both full-length Cx43 and GJA1-20k. Goat secondary antibodies conjugated to Alexa Fluor 647 and 555 (1:2500; Thermo Scientific) and imaged on a VersaDoc 5000 MP (Bio-Rad).

### Immunofluorescence

Cells were cultured on Nunc™ Lab-Tek™ II Chamber Slides™ (Thermo Fisher, 154526), fixed in 4% paraformaldehyde at 37°C for 20 min, and then stored in PBS at 4°C until staining. Cells were permeabilized with 0.2% Triton X-100 in PBS for 10 min then blocked with 5% goat serum in PBST (0.1% Tween-20 in PBS) for 1 hr. Rabbit anti-Cx43 (1:400, Sigma, C6219) was used as a primary antibody with a goat anti-rabbit secondary antibody conjugated to either Alexa Fluor 488 or 633 (1:500, Thermo Scientific). The anti-Cx43 antibody targets the C-terminal domain and will therefore identify both full-length Cx43 and GJA1-20k. Samples were mounted with cover glass using Pro-Long Glass Antifade Mountaint with NucBlue (Invitrogen, P36983) and imaged on a Zeiss LSM 710 Confocal Microscope using a 63x oil immersion objective.

### MSC and Chondrocyte Co-Cultures

*Contact Co-Cultures:* Chondrocytes with mCherry cytoplasm fluorescence were seeded onto 12-well plates for transfer quantification or on 2-well slides for imaging at a density of 35,000 and 40,000 cells, respectively, and cultured overnight in chondrocyte media (detailed above). MSCs with GFP mitochondrial fluorescence were added in a 1:2 MSC:chondrocyte ratio for transfer quantification or 1:10 ratio for imaging and co-cultured in 1X OptiMEM for 24 hrs. After 24 hrs, cells were processed for transfer quantification using flow cytometry (see below) or imaging. For imaging, cells were fixed in 4% PFA for 20 minutes, mounted with cover glass using the ProLong Glass Antifade Mountant with NucBlue Stain (Thermo Fisher, P36983), and imaged on a Zeiss LSM 710 Confocal Microscope using a 63x oil immersion objective. *Contact vs Transwell Co-Cultures:* Chondrocytes with GFP mitochondrial fluorescence were seeded on 12-well plates at a density of 100,000 cells/well and cultured overnight in chondrocyte media. MSCs with mCherry mitochondrial fluorescence were then seeded for co-cultures. For contact co-culture, 50,000 or 75,000 MSCs were added directly to the wells. For transwell co-cultures, 50,000 or 75,000 MSCs were seeded onto transwell inserts (1 μm pore size, Corning, 353103) and placed into wells containing chondrocytes. Both co-cultures were performed for 24 hrs in a 1:1 ratio of chondrocyte and MSC media and were then processed for flow cytometry to quantify transfer events.

### Mitochondrial Transfer Quantification

After co-culture, cells were rinsed with PBS, lifted, and fixed at 37°C in 4% PFA for 20 minutes in the dark. Cells were rinsed in PBS and re-suspended in 4°C FCM buffer (PBS with 0.5% BSA and 2mM EDTA) and placed in the fridge until analyzed. Cells were filtered (40 μm, Corning, 352235) and analyzed on a Thermo Fisher Attune NxT flow cytometer. Unstained cells and single-color control cells were used to set up gates for identifying transfer events. For contact co-cultures, quadrant gates were used to quantify MSC to chondrocyte transfer events as cells that had both mCherry (chondrocyte cytoplasm) and GFP (MSC mitochondria). For experiments comparing contact vs transwell co-cultures, an oval gate was used to identify MSC to chondrocyte mitochondrial transfer events (both GFP+ and mCherry+).

### Normalization and Correlation of GJA1 Expression and Mitochondrial Transfer

To investigate the relationship between MSC GJA1-43k (full length Cx43) and GJA1-20k expression with mitochondrial transfer, western blot data and mitochondrial transfer data were normalized to their respective controls. Specifically, the *GJA1* siRNA treated group was normalized to Stealth RNAi control, and the GJA1+ and GJA1-20k+ overexpression groups were normalized to the LacZ control group. Error propagation was calculated for each normalized value from standard deviations of the group. A linear correlation was performed with normalized GJA1-43k or GJA1-20k as the independent variable and the normalized percentage of mitochondrial transfer as the dependent variable.

### Statistics

Statistical analyses were performed using GraphPad Prism. Significant differences were analyzed by unpaired Student’s t-tests for comparisons between two groups. Mitochondrial transfer percentage was compared between transwell and contact co-culture groups using a two-way ANOVA. For all other comparisons between more than two groups, a one-way ANOVA was used.

For both one-way and two-way ANOVAs, a Tukey post-hoc test was performed for multiple comparisons. A p-value < 0.05 was considered significant. Significance is denoted with asterisks or with letters where groups not sharing a letter are significantly different.

## Results

### Oxidative stress increases incidence of MSC to chondrocyte mitochondrial transfer

To determine the effect of oxidative stress on mitochondrial transfer, chondrocytes were treated with t-BHP for 24 hrs (Fig 1Ai). Mitochondria are a major source of ROS, and t-BHP stimulation induced mitochondrial depolarization (indicated by JC-10 staining) and increased ROS production (indicated by CellROX green staining) in chondrocytes (Fig 1B-D). Chondrocyte mitochondria were depolarized in a dose-dependent manner with t-BHP stimulation where concentrations greater than 12 µM significantly increased the ratio of depolarized to polarized mitochondria compared to unstimulated controls (p<0.05, Fig 1B). Similarly, 12 µM and 30 µM concentrations of t-BHP increased ROS in chondrocytes compared to unstimulated controls (p<0.05, Fig 1CD). Chondrocytes with mCherry cytoplasmic fluorescence were co-cultured with MSCs that had GFP mitochondrial fluorescence in 2D contact co-cultures for 24 hrs (Fig 1A). Confocal imaging confirmed mitochondrial transfer events with Z-stacks identifying the presence of GFP fluorescence within chondrocyte cell bodies (Fig 1E). Flow cytometry was used to quantify the percentage of mitochondrial transfer events. A quadrant gate with single color controls was used to identify events both mCherry+ and GFP+ as transfer events (Fig 1F). Chondrocytes with t-BHP-induced oxidative stress had significantly more mitochondrial transfer events compared to unstimulated chondrocytes (p<0.001, Fig 1G-I).

**Figure 1.**
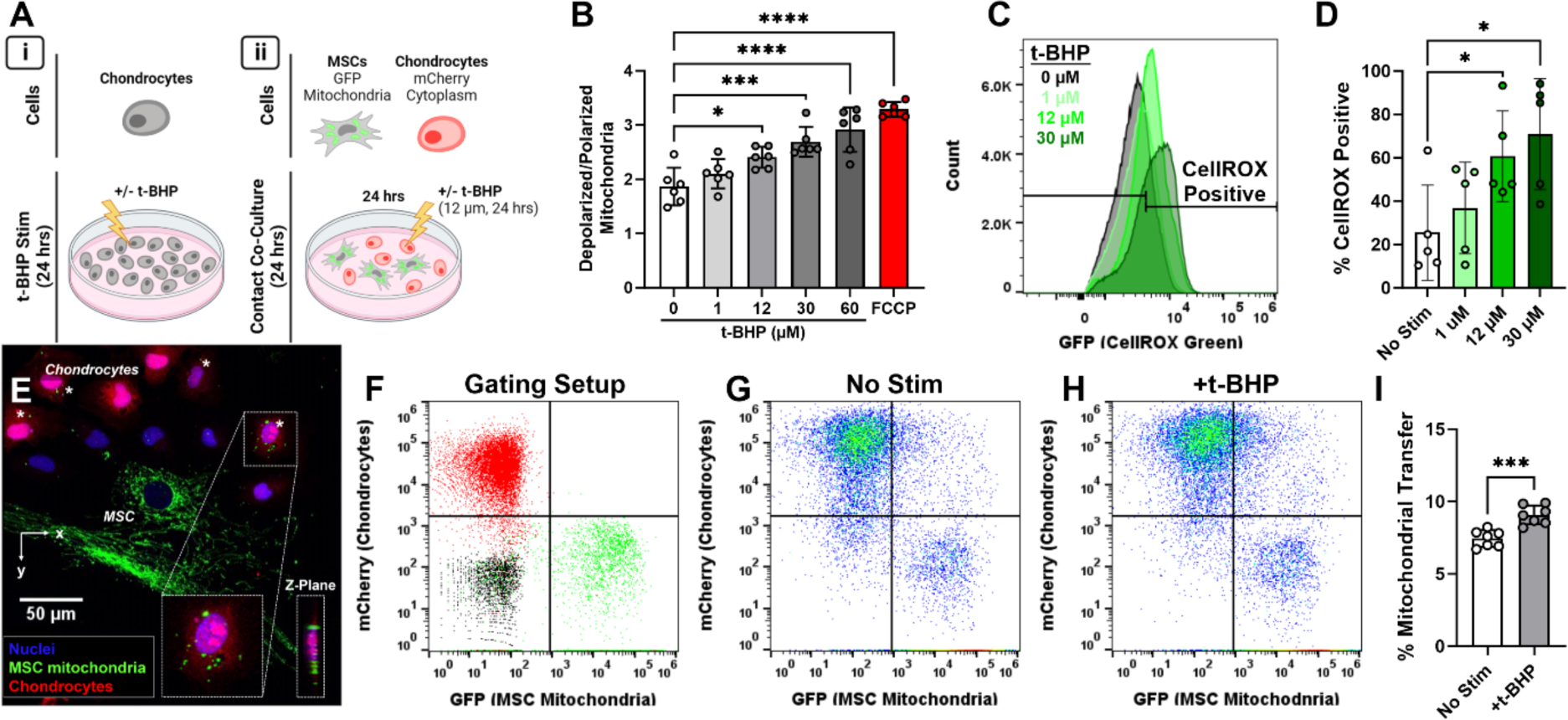
t-BHP-induced oxidative stress in chondrocytes increases incidence of mitochondrial transfer from MSCs. (A) Chondrocytes were cultured alone with or without t-BHP (12 µM, 24 hrs) to quantify mitochondrial polarization and ROS production (i) and prior to co-culture with MSCs (ii). (B) Ratio of depolarized to polarized mitochondria in chondrocytes after t-BHP or FCCP stimulation (n=5-6). (C) Histograms represent ROS quantification for chondrocytes stimulated with t-BHP (pooled across n=5 replicates). (D) Quantification of percentage of chondrocytes that were positive for CellROX green staining on flow cytometry (n=5). (E) Z-projection of co-culture with chondrocytes and MSCs (red: chondrocytes; green: MSC mitochondria; blue: nuclei). Quadrant gate showing representative flow data for single color controls (F), co-cultures with unstimulated chondrocytes (G), and co-cultures with t-BHP stimulated chondrocytes (H). (I) Quantification of mitochondrial transfer from co-cultures (n=7). *p<0.05, **p<0.01, ***p<0.001, ****p<0.0001. Created with BioRender.com.

*MSC-chondrocyte mitochondrial transfer occurs predominantly through actin cellular extensions* To determine the importance of direct cell contacts in mitochondrial transfer between MSCs and chondrocytes, co-cultures were performed for 24 hrs in direct contact (cell culture plates) or without direct contact (transwell inserts, Fig 2A). For these experiments, both cell types had lentiviral-labeled mitochondria (chondrocytes: GFP mitochondria; MSCs: mCherry mitochondria). After 24 hrs of co-culture, contact co-cultures showed many transfer events (mCherry+ and GFP+) whereas the transwell co-cultures had few transfer events (Fig 2B-D). Notably, the transwell co-cultures had minimal events as only mCherry+, indicating the MSCs did not migrate across the transwell insert (Fig 2D). Contact co-cultures with a higher MSC seeding density had a higher incidence of mitochondrial transfer (p<0.01), but there was no difference in transfer between transwell co-culture groups with 50k vs 75k MSC seeding density (p=0.99). Contact co-cultures had significantly higher incidence of transfer events compared to transwell co-cultures for both MSC seeding densities (p<0.0001).

**Figure 2.**
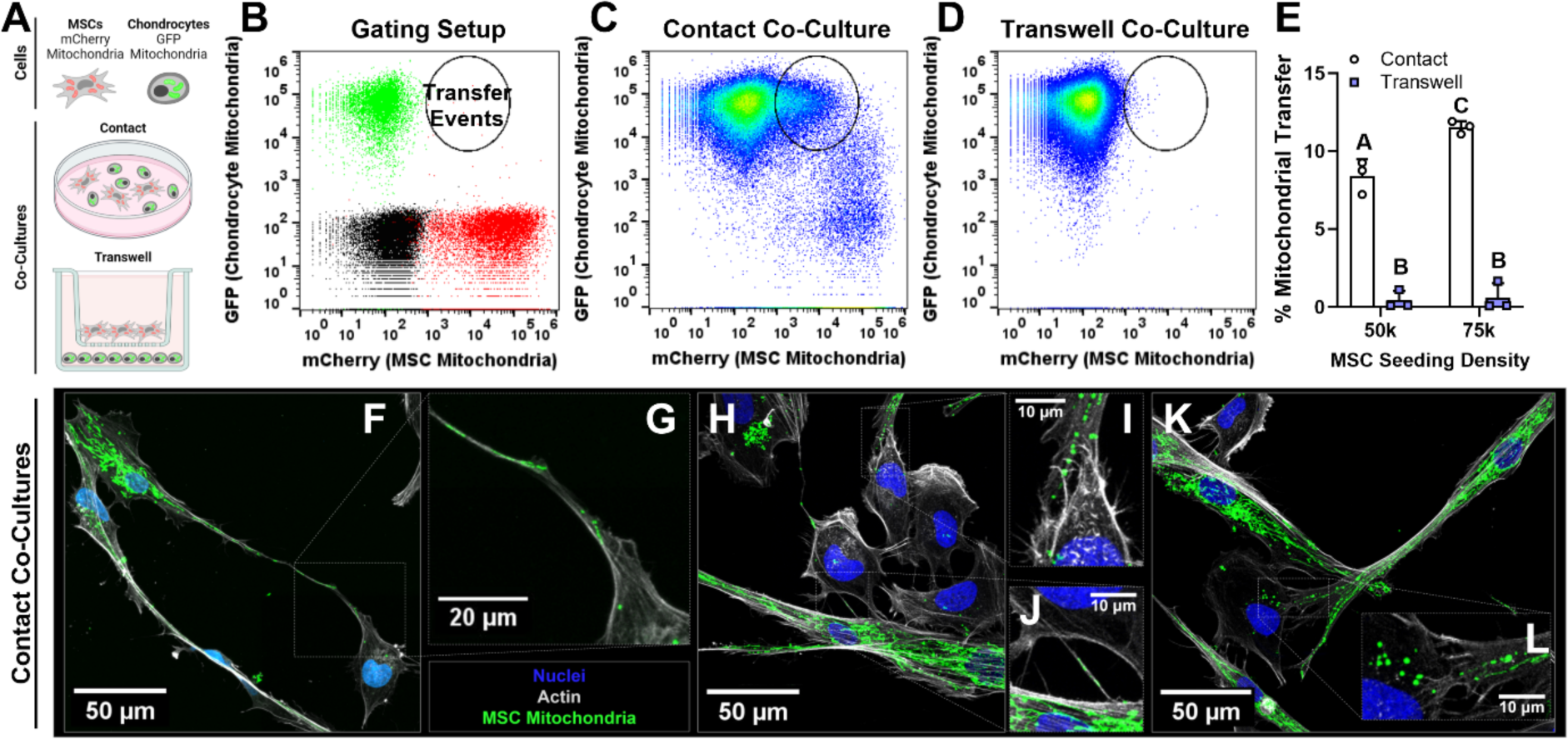
Direct cell-cell contacts are critical for MSC-Chondrocyte mitochondrial transfer. (A) Chondrocytes (GFP mitochondrial fluorescence) and MSCs (mCherry mitochondrial fluorescence) were co-cultured for 24 hrs in either 2D contact on cell culture plates or with no contact using transwell inserts. (B) Single color controls were used to establish gating strategy for identifying transfer events. Representative data for co-cultures from contact (C) and transwell (D) co-cultures. (E) Quantification of mitochondrial transfer from flow cytometry (n=3, groups not sharing a letter are significantly different, p<0.01). (F-L) Z-projections from 2D contact co-cultures captured using confocal imaging with MSCs (GFP mitochondria) and unlabeled chondrocytes (green: MSC mitochondria; grey: actin; blue: nuclei). Created with BioRender.com.

To examine cell-cell interactions, cells were fixed after 24 hrs of contact co-culture and stained with phalloidin to visualize the F-actin cytoskeleton for both cell types. For these imaging experiments, MSCs with GFP mitochondria and chondrocytes with no fluorescence were used. Confocal imaging revealed MSC actin cellular extensions in contact with adjacent chondrocytes that contained green fluorescence, indicating mitochondrial transfer had occurred (Fig 2F-L). Notably, MSC mitochondria were located within these actin cellular extensions (Fig 2G,J).

### Cx43 staining is enriched along actin cellular extensions containing mitochondria in MSCs

To investigate the role of Cx43 in mitochondrial transfer between MSCs and chondrocytes, Cx43 immunofluorescence was performed after 24 hrs of contact co-culture. MSC actin extensions that contained mitochondria also had enriched Cx43 staining at the end of the actin cell processes (Fig 3). Actin cell extensions ranged in size and measured over 150 µm in length from the cell body (Fig 3B). Notably, the Cx43 staining at the actin cellular extensions was more diffuse and less punctate, which is indicative of the truncated Cx43 isoforms and not hemichannels or gap junction plaques (36).

**Figure 3.**
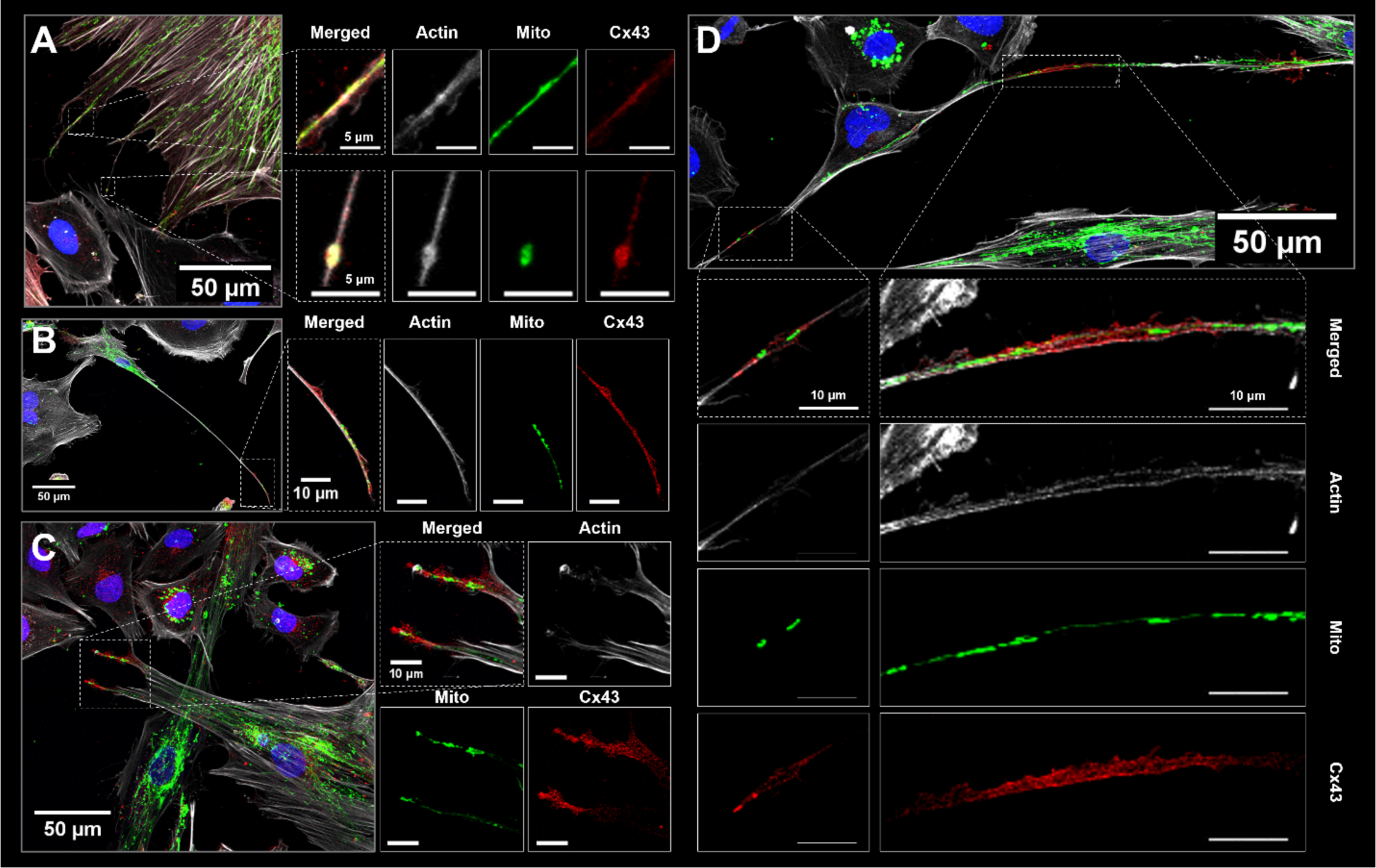
Cx43 staining is enriched along actin cellular extensions containing mitochondria in MSCs. MSC actin cellular extensions were observed in contact co-cultures with chondrocytes after 24 hrs. MSCs with GFP mitochondria and unstained chondrocytes were used in these experiments. Cells were stained for Cx43 and actin (phalloidin). Merged z-projections and individual channels are shown (green: MSC mitochondria; red: Cx43; grey: actin; blue: nuclei).

### Knocking-down Cx43 expression in MSCs decreases mitochondrial transfer to chondrocytes

To evaluate the role of Cx43 expression in mediating mitochondrial transfer between MSCs and chondrocytes, siRNA was used to knock-down Cx43 expression in MSCs prior to co-culture. Knock-down of Cx43 in MSCs was confirmed using western blot and normalized to α-tubulin (Fig 4A). Cx43 was detected using an antibody targeting the C-terminus of the protein and therefore can detect both full-length Cx43 (43k) and truncated isoforms. Quantification showed a significant decrease in full-length Cx43 and the truncated isoform GJA1-20k (n=3; p<0.0001 and p<0.05, respectively). Interestingly, *GJA1* siRNA increased the relative expression of the 20k isoform to the full-length protein (GJA1-20k/Cx43, p<0.01). As both Cx43 and GJA1-20k arise from the same mRNA, this may be due to GJA1-20k potentially having a longer half-life. Additionally, immunofluorescence confirmed the knock-down of Cx43 in MSCs where there was a decrease in the punctate and diffuse staining within the cells (Fig 4E,F). *GJA1* siRNA treated and control (Stealth RNAi treated) MSCs with GFP mitochondrial fluorescence were co-cultured with chondrocytes for 24 hrs (chondrocytes with mCherry cytoplasmic fluorescence were used for these experiments). Chondrocytes co-cultured with *GJA1* siRNA treated MSCs had significantly less mitochondrial transfer events compared to control MSCs (p<0.01, Fig 4G-I). Notably, knocking down Cx43 in the MSCs did not completely suppress mitochondrial transfer to chondrocytes.

**Figure 4.**
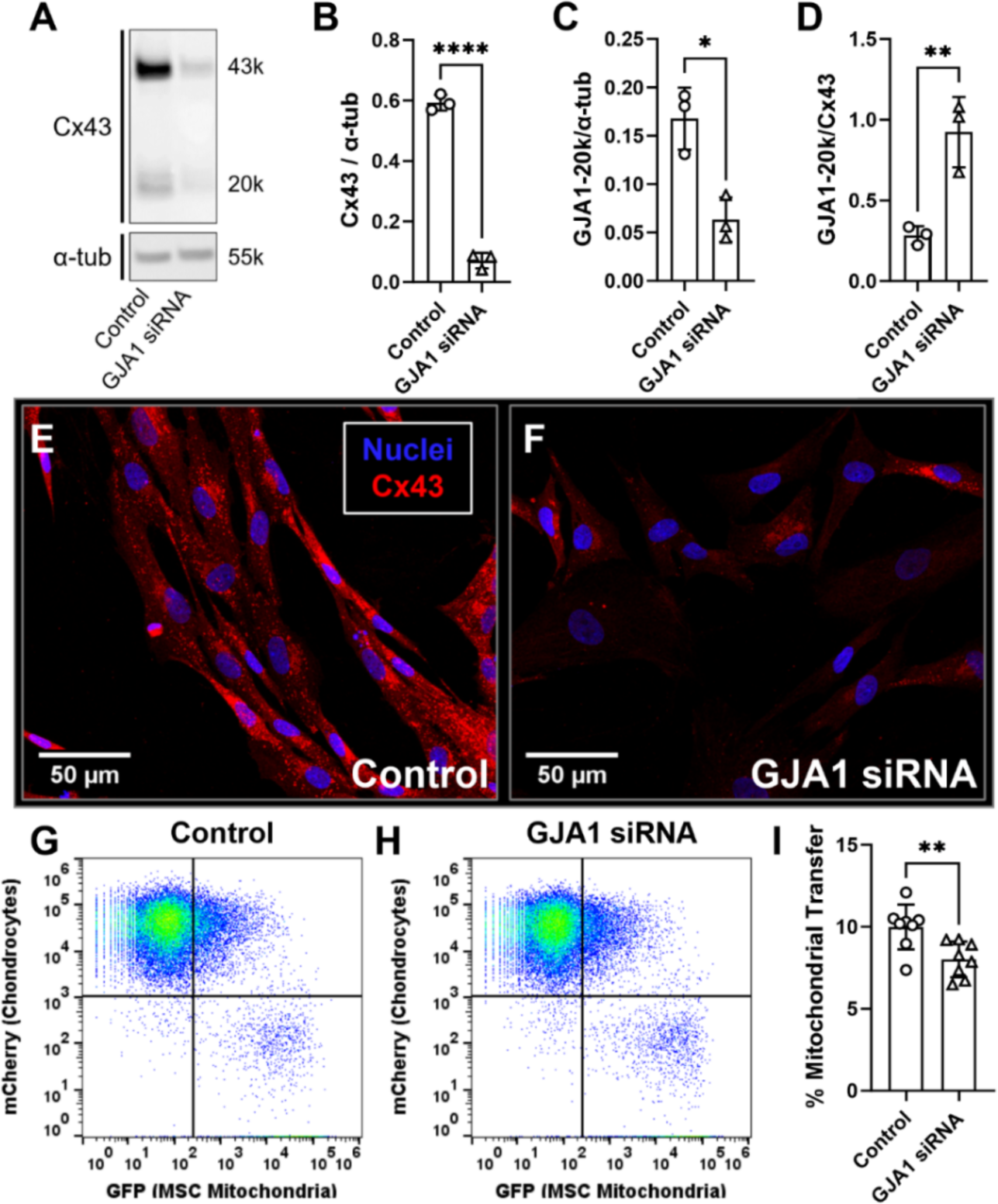
GJA1 siRNA knocks-down Cx43 expression in MSCs and decreases incidence of mitochondrial transfer to chondrocytes. (A) Western blot analysis of full length Cx43 (GJA1-43k) and the truncated isoform GJA1-20k in MSCs after GJA1 siRNA treatment. Cx43 (B) and GJA1-20k (C) expression were normalized to α-tubulin (n=3). (D) Relative isoform expression was calculated as the ratio of GJA1-20k to Cx43 (n=3). (E-F) Representative images of Cx43 immunofluorescence for control (Stealth RNAi) and GJA1 siRNA treated MSCs. (G-H) Representative data from flow cytometry analyses of co-cultures. (I) Quantification of mitochondrial transfer events from co-cultures using flow cytometry (n=8). Full-length western blot for quantification is presented in Supplementary Figure 2. *p<0.05, **p<0.01, ***p<0.001, ****p<0.0001.

### MSC GJA1-20k expression mediates mitochondrial transfer to chondrocytes

To investigate the roles of full-length Cx43 (GJA1-43k) and GJA1-20k in mediating mitochondrial transfer from MSCs to chondrocytes, lentiviral transduction was performed to overexpress Cx43 (*GJA1*+, all isoforms) or GJA1-20k (GJA1-20k+) in MSCs. Overexpression of Cx43 and GJA1-20k was confirmed using western blot and immunofluorescence (Fig 5A-G). Western blot quantification confirmed a significant increase in full-length Cx43 in GJA1+ MSCs compared to LacZ (control) MSCs (p<0.05), and a significant increase in GJA1-20k in GJA1+ and GJA1-20k+ MSCs compared to LacZ (control) MSCs (p<0.05 and p<0.01, respectively). The ratio of GJA1-20k to full-length Cx43 was significantly higher in the GJA1-20k+ MSCs compared to both *GJA1*+ and LacZ MSCs, indicating an increased expression of the 20k isoform relative to the full-length Cx43 expression (p<0.01). *GJA1*+ MSCs had enriched diffuse and punctate Cx43 staining compared to LacZ MSCs. Notably, GJA1-20k+ MSCs had enriched diffuse Cx43 staining throughout the cell body compared to both GJA1+ and LacZ MSCs but no apparent difference in punctate Cx43 staining compared to LacZ (Fig 5E-G). This is consistent with prior work demonstrating localization of GJA1-20k to the cytoplasm within the vesicular transport pathway and mitochondria (36,40,44).

**Figure 5.**
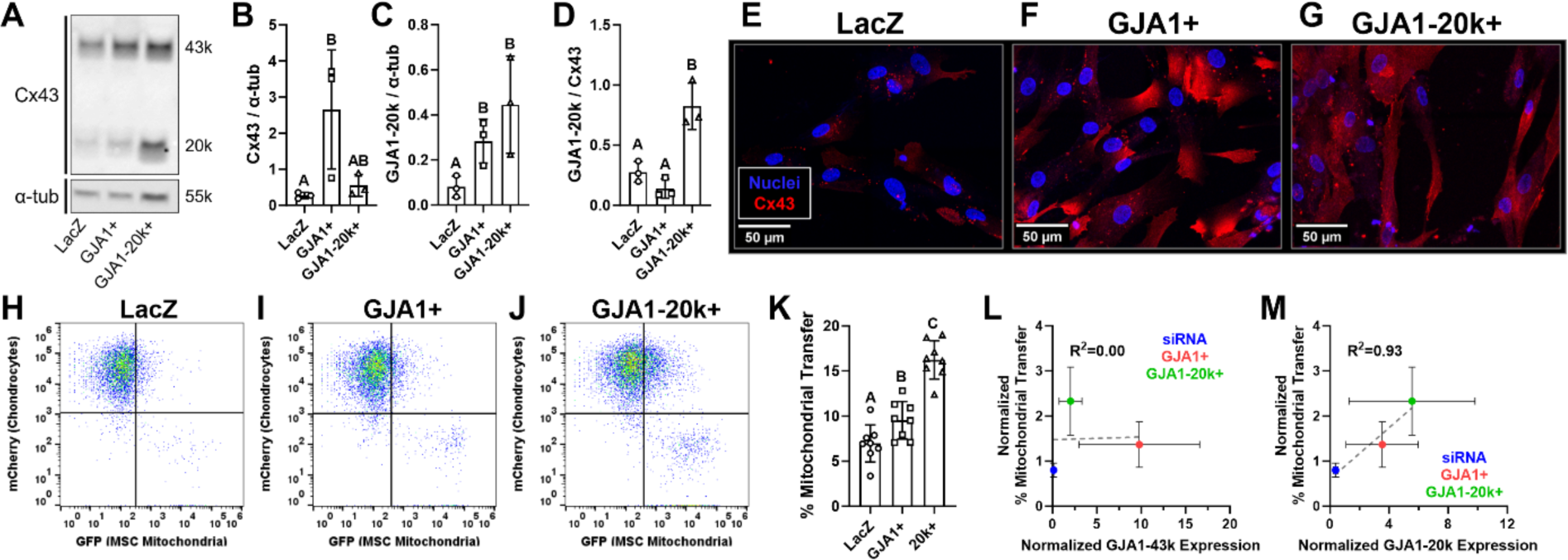
Overexpression of GJA1-20k in MSCs increases mitochondrial transfer to chondrocytes and MSC GJA1-20k expression correlates with incidence of mitochondrial transfer. Western blot analysis of full length Cx43 (GJA1-43k) and the truncated isoform GJA1-20k in MSCs after lentiviral transduction (LacZ (control), GJA1+, GJA1-20k+). Cx43 (B) and GJA1-20k (C) expression were normalized to α-tubulin (n=3). (D) Relative isoform expression was calculated as the ratio of GJA1-20k to Cx43 (n=3). (E-G) Representative images of Cx43 immunofluorescence for LacZ (control), GJA1+, and GJA1-20k+ MSCs. (H-J) Representative data from flow cytometry analyses of co-cultures. (K) Quantification of mitochondrial transfer events from co-cultures using flow cytometry (n=8). (L-M) Linear correlations for normalized protein expression of full-length Cx43 (GJA1-43k) and GJA1-20k with normalized incidence of mitochondrial transfer (groups normalized to their respective controls). Groups not sharing a letter are significantly different (p<0.05). Full-length western blot for quantification is presented in Supplementary Figure 3.

GJA1+ and GJA1-20k+ MSCs were co-cultured with chondrocytes for 24 hrs and evaluated using flow cytometry to quantify the incidence of mitochondrial transfer. GJA1+ MSCs had an increased incidence of mitochondrial transfer compared to LacZ controls (p<0.05). Interestingly, GJA1-20k+ MSCs had the highest incidence of mitochondrial transfer that was greater than co-cultures with GJA1+ MSCs and LacZ control MSCs (p<0.001 and p<0.0001, respectively; Fig 5K). To further understand the relationship between Cx43 isoform expression and incidence of mitochondrial transfer from MSCs to chondrocytes, linear correlations were performed with normalized protein expression and normalized mitochondrial transfer values. These analyses show no correlation between full-length Cx43 expression in MSCs and the incidence of mitochondrial transfer to chondrocytes (R^2^=0.00, Fig 5L). In contrast, there is a strong correlation between the normalized expression of GJA1-20k in MSCs and the incidence of mitochondrial transfer to chondrocytes (R^2^=0.93, Fig 5M).

## Discussion

The objectives of this study were to investigate the role of oxidative stress on MSC to chondrocyte mitochondrial transfer and to examine the role of Cx43 in the mechanism of transfer. The data from these studies show that t-BHP-induced oxidative stress increases mitochondrial transfer from MSCs to chondrocytes. Additionally, our findings support the hypothesis that GJA1-20k is a key mediator of mitochondrial transfer through direct cell-cell contacts.

While there is promising evidence that MSC injections preserve cartilage tissue after injury in the short-term, the potential mechanisms for this therapeutic effect are poorly understood. Mitochondrial transfer could be a driving cause of the observed therapeutic benefit, as the transfer of mitochondria from MSCs to stressed cells has been shown to be protective against injury *in vivo* in lung epithelial and tendon cells (13,24). Additionally, mitochondrial transfer has been shown to increase when cells are stressed both *in vitro* and *in vivo* (13,25). Here, we found similar results, whereby inducing oxidative stress with t-BHP in chondrocytes increased the incidence of mitochondrial transfer from MSCs.

Cx43 has been implicated in mitochondrial transfer through multiple mechanisms including gap junction communication (13,25,31) and gap junction internalization (33). Gap junction signaling through Cx43 was found to stimulate the formation of TNTs and microvesicles in MSCs (13), which are two mechanisms of mitochondrial transfer that have been identified for MSCs to chondrocytes (25,45). Recently, over-expression of the truncated isoform GJA1-20k in astrocytes was found to significantly increase mitochondrial transfer to neurons *in vitro* (39). GJA1-20k has been identified as a regulator of mitochondrial motility and dynamics through interactions with microtubules and the actin cytoskeleton (37,43,44,46), but to the authors’ knowledge has not previously been investigated in MSCs. Here, we report that MSCs express GJA1-20k and that MSC expression of GJA1-20k strongly correlates with the incidence of mitochondrial transfer. These data suggest that GJA1-20k aids in the transport of mitochondria from MSCs to adjacent cells and may assist through mobilization of mitochondria through TNTs, packaging of mitochondria within microvesicles (EVs), or through the formation of actin cellular extensions (Fig 6). Transwell co-cultures resulted in significantly lower levels of mitochondrial transfer compared to contact co-cultures, suggesting direct cell-contacts are the primary route of transfer. Microvesicle-meditaed mitochondrial transfer can occur from MSCs to chondrocyte but has been reported at low efficiencies (<1%) (45). While there are likely multiple mechanisms of mitochondrial transfer occurring between MSCs and chondrocytes, these data highlight the role of GJA1-20k in increasing transfer events.

**Figure 6.**
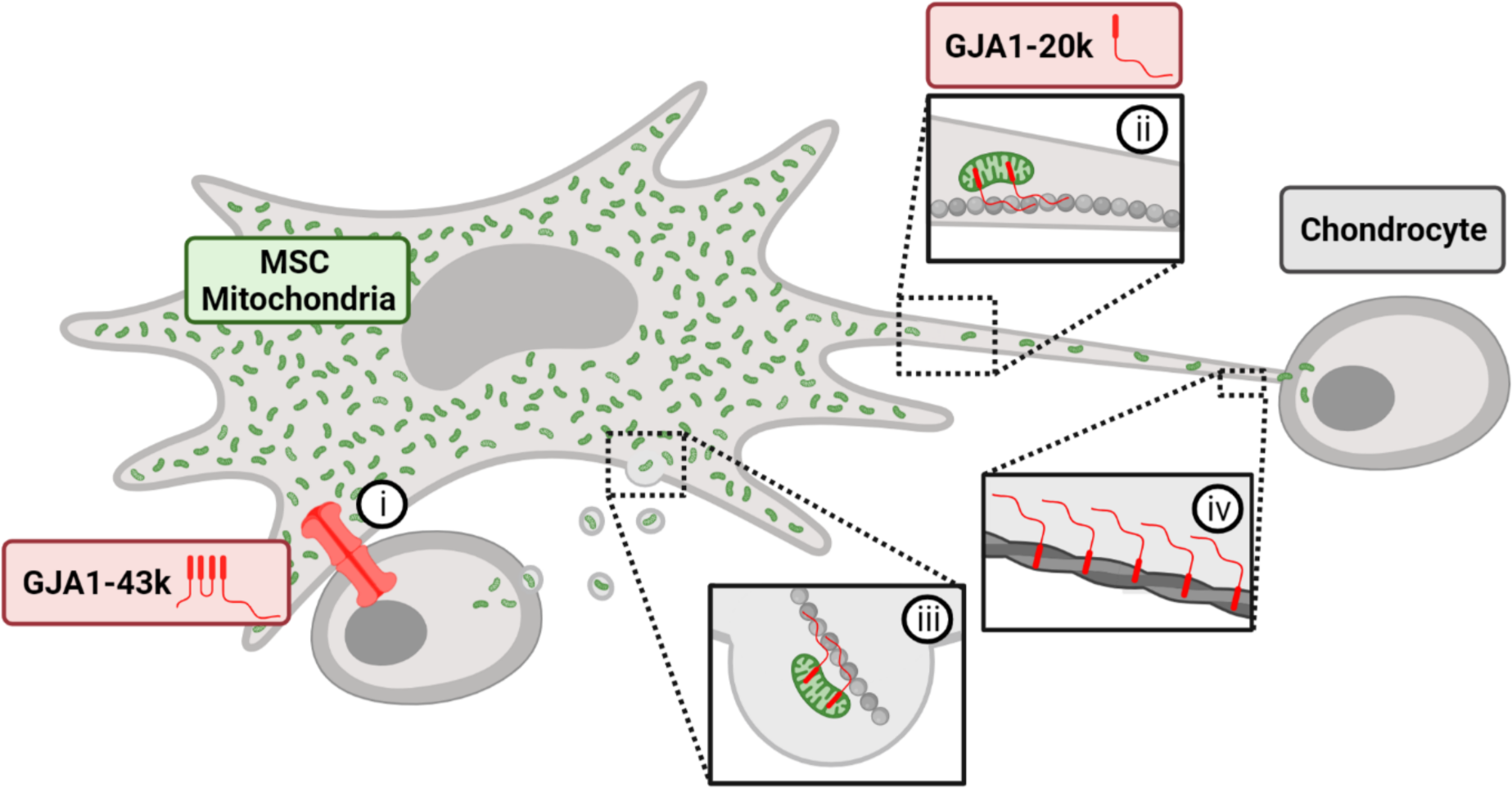
Hypothesized role of Cx43 and GJA1-20k in MSC to chondrocyte mitochondrial transfer. Previous work has identified the role of gap junction communication in MSC to chondrocyte mitochondrial transfer (i) (Fahey+ *Scientific Reports* 2022). Further, the results from this study suggest that GJA1-20k aids in the transport of mitochondria from MSCs to adjacent cells and may assist through mobilization of mitochondria through TNTs (ii), packaging of mitochondria within microvesicles (iii), or through the formation of actin cellular extensions (iv). Created with BioRender.com.

Here, ectopic alteration of Cx43 expression in the MSC population caused significant changes in the percentage of mitochondrial transfer; whereby knocking down Cx43 decreased transfer events and overexpression of Cx43 and GJA1-20k increased transfer events. While these data suggest the importance of Cx43 in transferring mitochondria from MSCs to chondrocytes, the Cx43 expression of chondrocytes could also influence mitochondrial transfer between the cells. Chondrocytes have increased expression of Cx43 in OA cartilage tissue (47), and this could increase the rate of mitochondrial transfer from MSCs as increased Cx43 gap junction communication was found to increase TNT and microvesicle release from MSCs (13). Therefore, overexpression of Cx43 in chondrocytes, like that seen in OA, may lead to increased mitochondrial transfer events from MSCs. t-BHP was used in this study to induce oxidative stress and resulted in an increase in transfer events but did not affect Cx43 or GJA1-20k expression in chondrocytes (Supplementary Figure 1). These data suggest oxidative stress in chondrocytes can drive increased mitochondrial transfer through pathways independent of changes in Cx43 protein expression.

Altering the expression of Cx43 in the MSC population significantly changed the incidence of mitochondrial transfer between cells. GJA1 siRNA was used to knockdown protein expression of both full-length (GJA1-43k) and the truncated isoform GJA1. Knocking down Cx43 in MSCs significantly decreased mitochondrial transfer but did not fully suppress transfer from MSCs to chondrocytes. Notably, overexpressing the GJA1-20k isoform in MSCs resulted in the highest increase in mitochondrial transfer to chondrocytes compared to overexpression of LacZ or GJA1+. GJA1-20k is critical for trafficking Cx43 hemichannels from the Golgi apparatus to the cell membrane where gap junctions can form (36,40). Expression of GJA1-20k is regulated by multiple cell signaling pathways including Smad3, ERK, P13K/Akt/mTOR, and MNK1/2, and has been implicated as a means for cells to control Cx43 hemichannel and gap junction formations post-transcriptionally (36,40,48,49). In addition to the role GJA1-20k may play in mitochondrial transfer through the actin cytoskeleton and trafficking of mitochondria, elevated levels of GJA1-20k may also increase Cx43 hemichannels and gap junctions at the MSC cell membrane. Previous reports have identified the importance of Cx43 channel function in mitochondrial transfer (13,25,31), and therefore elevated levels of GJA1-20k may increase transfer through augmented trafficking of GJA1-43k hemichannels to the cell membrane.

In this study, mitochondrial transfer from MSCs to chondrocytes was evaluated in monolayer cell culture which does not account for the dense ECM chondrocytes are surrounded by *in vivo*. Recent work has confirmed that MSCs can transfer mitochondria to chondrocytes in situ (25), highlighting the potential therapy of mitochondrial transfer from MSCs to chondrocytes in their native tissue environment. The results presented here further our understanding of the mechanisms of mitochondrial transfer from MSCs to chondrocytes, which can inform the development of MSC-based therapies for treating cartilage defects and OA. There is conflicting evidence about the role of Cx43 gap junctions in mitochondrial transfer as some studies that chemically inhibited gap junctions (carboxonolone disodium) and Cx43 channel (Gap26, Gap27) functions found a decrease in mitochondrial transfer (13,25), yet one study found chemical inhibition (Gap26) had no effect on the incidence of mitochondrial transfer (50). This discrepancy could be explained by the differences in cell types evaluated in the studies as multiple mechanisms of mitochondrial transfer have been described across different cell populations (21). In addition to Cx43, other proteins have been identified as regulators of mitochondrial transfer between MSCs and other cell types including Miro1 (19,21,22). While multiple proteins are likely involved in regulating mitochondrial transfer, the data presented here identified Cx43 as a critical mediator of transfer between MSCs and chondrocytes. Additionally, our work incorporates the impact of GJA1-20k to reveal a key role for this internally translated Cx43 isoform, for the first time, in mitochondrial transfer between MSCs and chondrocytes.

## Conclusions

The data presented in this study highlight a role for the truncated Cx43 isoform GJA1-20k in MSC to chondrocyte mitochondrial transfer and is the first to evaluate the role of GJA1-20k in MSC mitochondrial transfer. Our results identify direct cell-cell contacts as the primary route of mitochondrial transfer, with GJA1-20k serving as a key mediator in this process. Collectively, these data highlight the potential of GJA1-20k as a therapeutic target for promoting mitochondrial transfer as a regenerative therapy for cartilage tissue repair in OA.

## List of Abbreviations

Cx43: Connexin 43
DMEM: Dulbecco’s modified Eagle’s medium
DTT: Dithiothreitol
ECM: Extracellular matrix
FCCP: Carbonyl cyanide p-trifluoro-methoxyphenyl hydrazone
GFP: Green fluorescence protein
GJA1: Gap junction protein alpha 1 (gene name for Cx43)
GJA1-43k: Full length 43k isoform of GJA1
GJA1-20k: Truncated 20k isoform of GJA1
LacZ: Gene encoding β-galactosidase
MSC: Mesenchymal stromal cell
OA: Osteoarthritis
PBS: Phosphate-buffered saline
PFA: Paraformaldehyde
RNA: Ribonucleic acid
ROS: Reactive oxygen species
siRNA: Short interfering RNA
t-BHP: tert-Butyl hydroperoxide
TNT: Tunneling nanotubules

## Declarations

### Ethics approval and consent to participate

This manuscript includes the use of an immortalized human chondrocyte line and human MSCs that are commercially available. As such, the need for ethical approval for procurement of the cell lines does not apply to these studies.

### Consent for publication

Not applicable.

### Availability of data and materials

The datasets used and/or analyzed during the current study are available from the corresponding author on reasonable request.

### Author’s contributions

RMI contributed to the study design, data collection, data analysis, data interpretation, and drafting of the paper. MAT and MJF contributed to data collection, data analysis, data interpretation, and editing of the paper. MDM, JWS, and MLD participated in the study design, data interpretation, and editing of the paper. All authors have read and approved the current version of the manuscript.

## Acknowledgements

Imaging and flow cytometry data were acquired through the Cornell Institute of Biotechnology’s BRC Imaging and Flow Cytometry Facilities (RRID:SCR_021741 and RRID:SCR_021740, respectively).

---

**Supplemental Figure 1.**
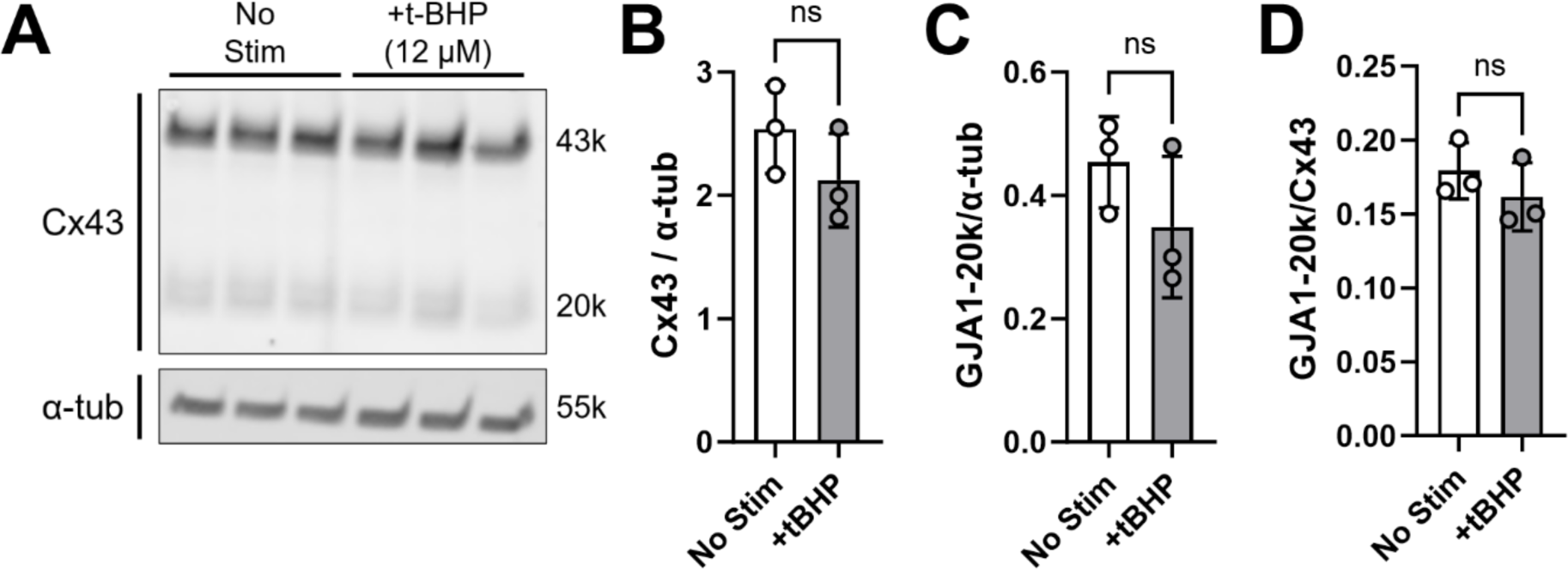
t-BHP does not alter Cx43 expression in chondrocytes. Chondrocytes were cultured for 24 hrs with or without t-BHP (12 μM) and lysed for western blotting (A). There was no significant difference between unstimulated chondrocytes and the t-BHP stimulated chondrocytes for normalized Cx43 expression (B), normalized GJA1-20k expression (C), or the ratio of GJA1-20k to Cx43 expression (D).

**Supplementary Figure 2.**
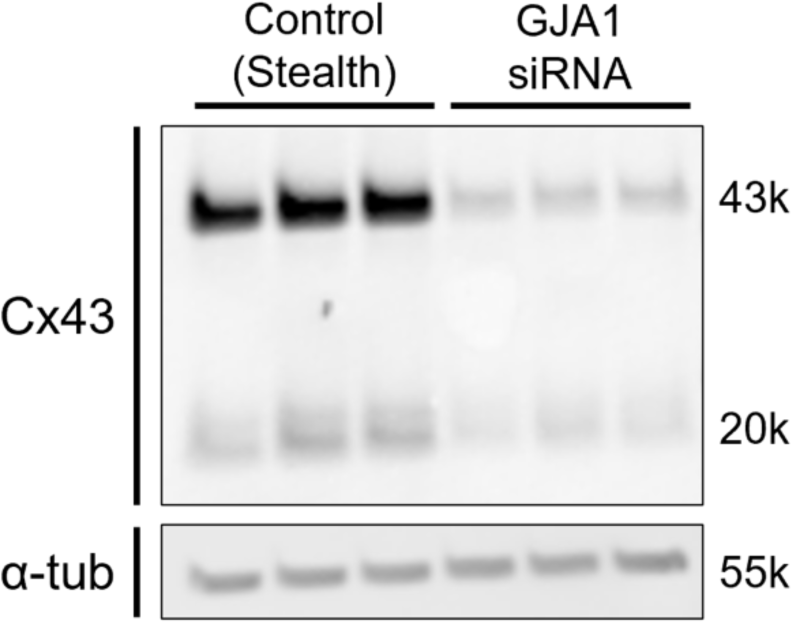
Western blot for GJA1 siRNA quantification. Western blot used for quantification of Cx43 normalized to α-tubulin presented in Figure 4. Additionally, a cropped version of this western blot is shown in Figure 4.

**Supplementary Figure 3.**
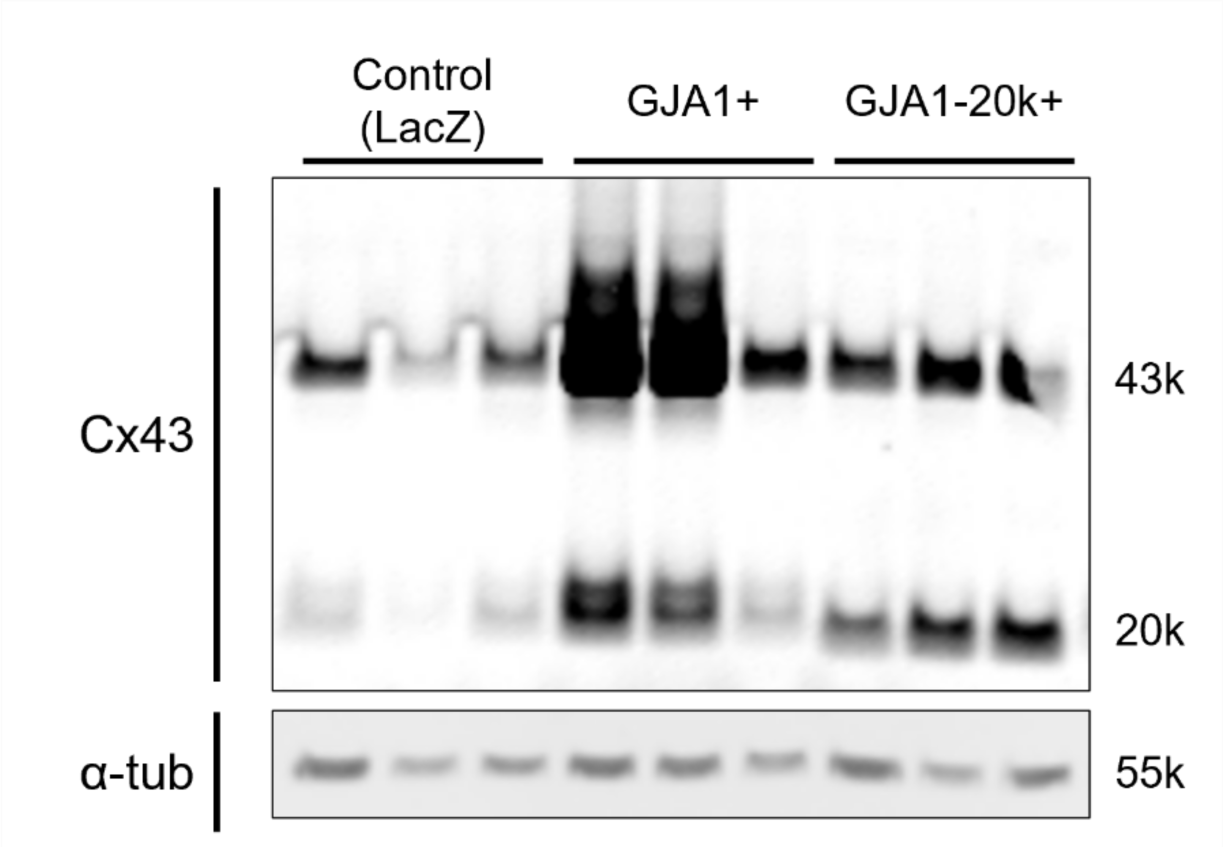
Western blot for GJA1+ and GJA1-20k+ overexpression quantification. Western blot used for quantification of Cx43 normalized to α-tubulin presented in Figure 5.

Supplemental File 4. Original Western blot images.

**Figure 1.**
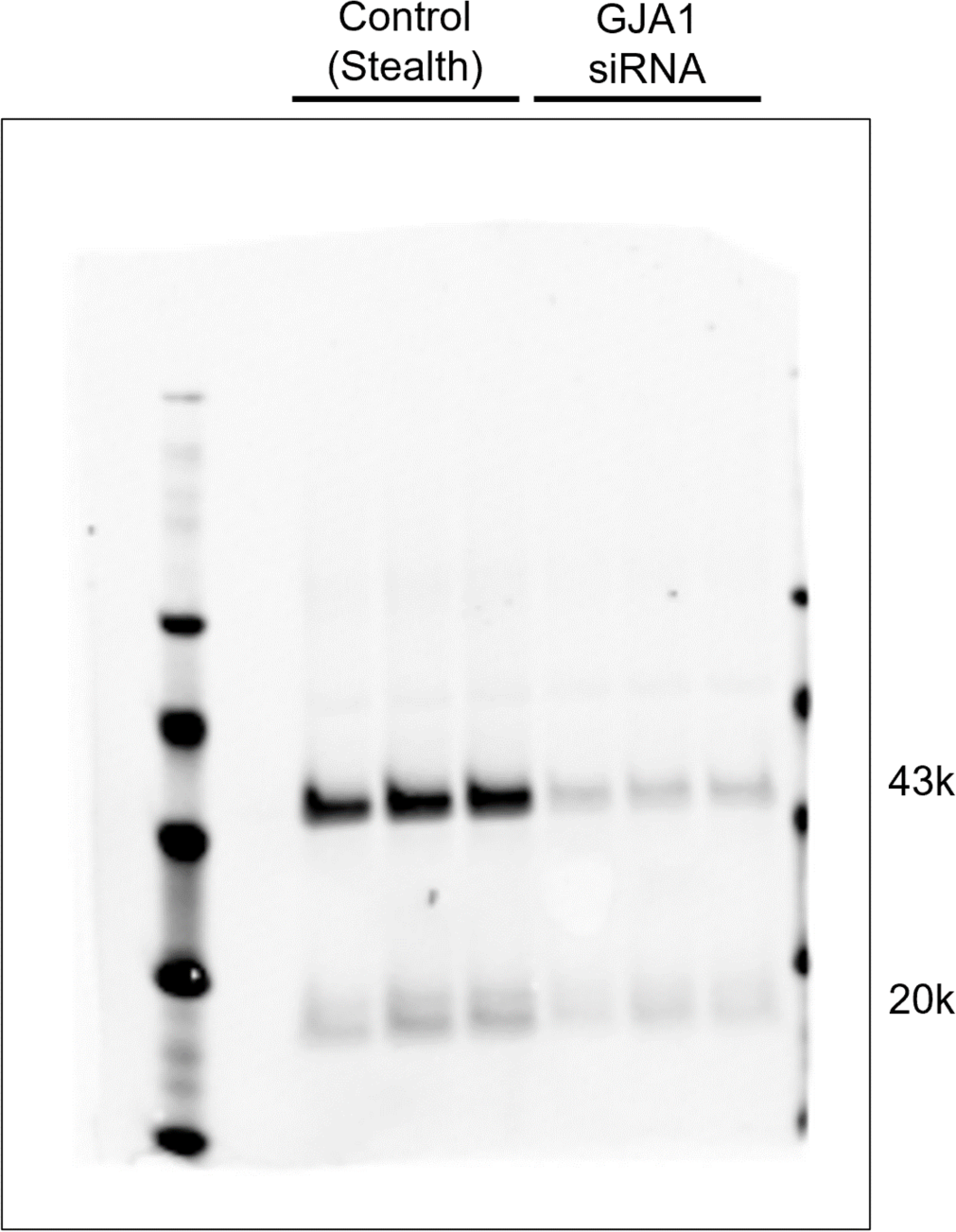
Western blot for siRNA treated MSCs: Cx43 quantification.

**Figure 2.**
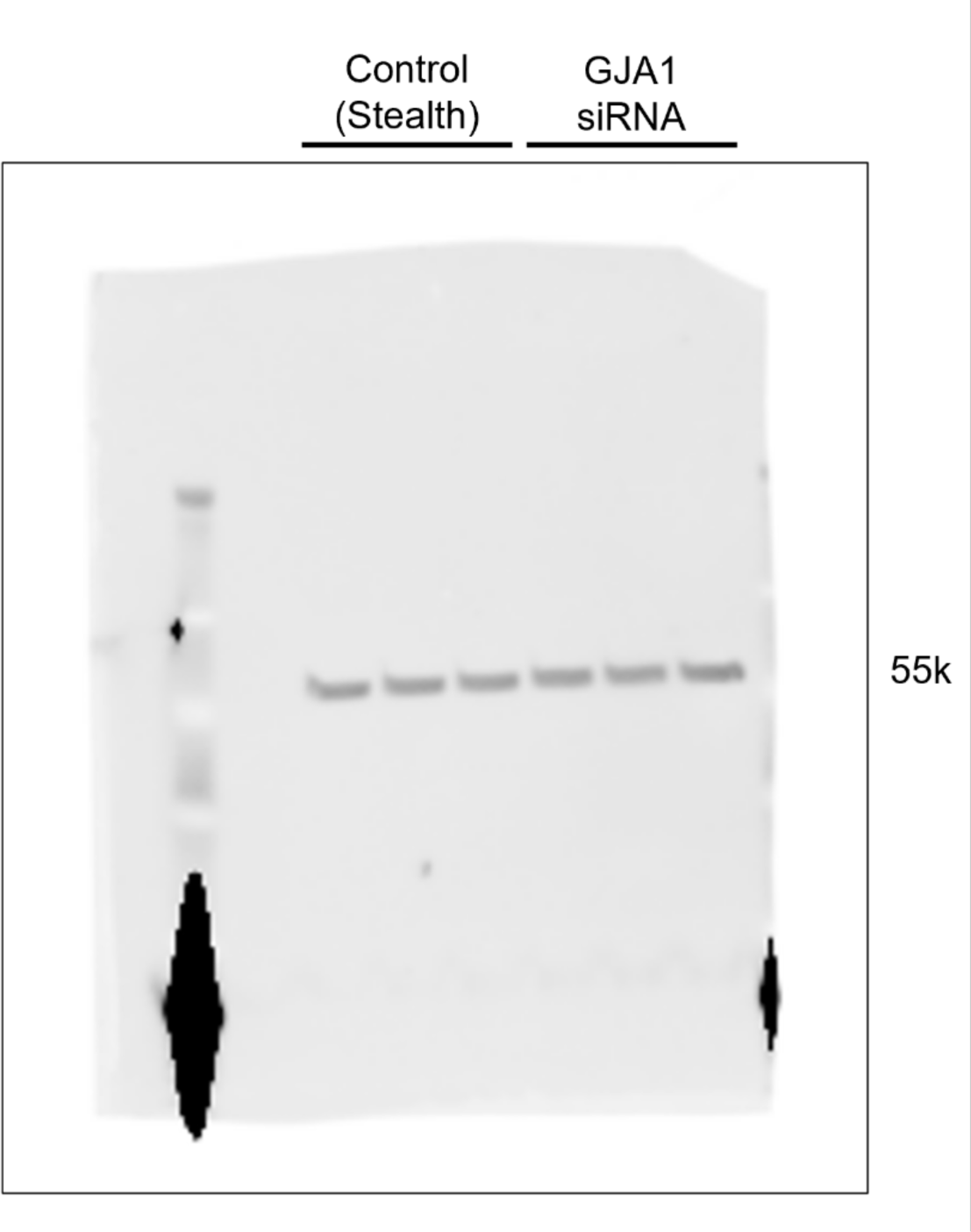
Western blot for siRNA treated MSCs: α-tubulin quantification.

**Figure 3.**
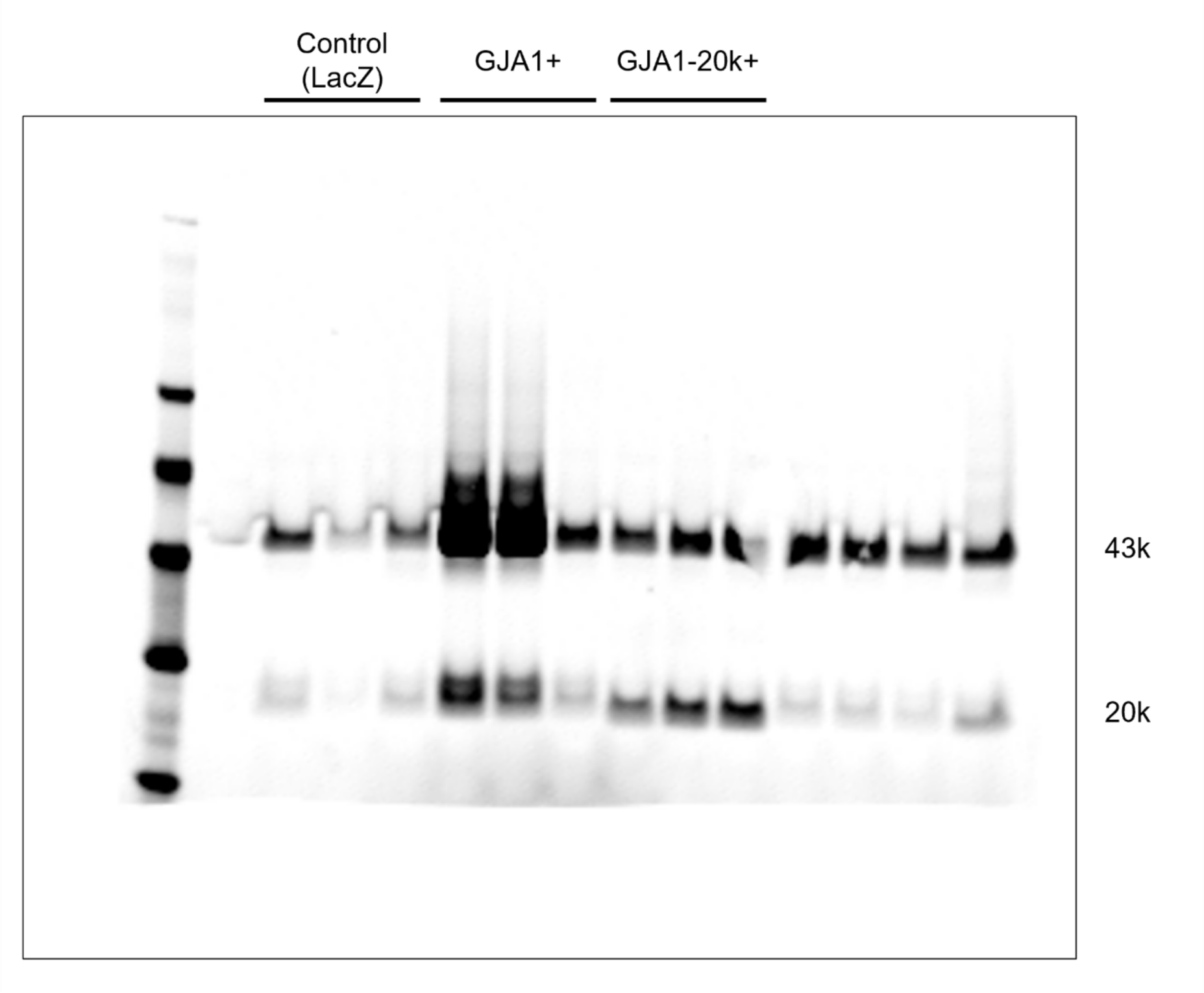
Western blot for LacZ, GJA1+, and GJA1-20k+ lentiviral transduced MSCs: Cx43 quantification.

**Figure 4.**
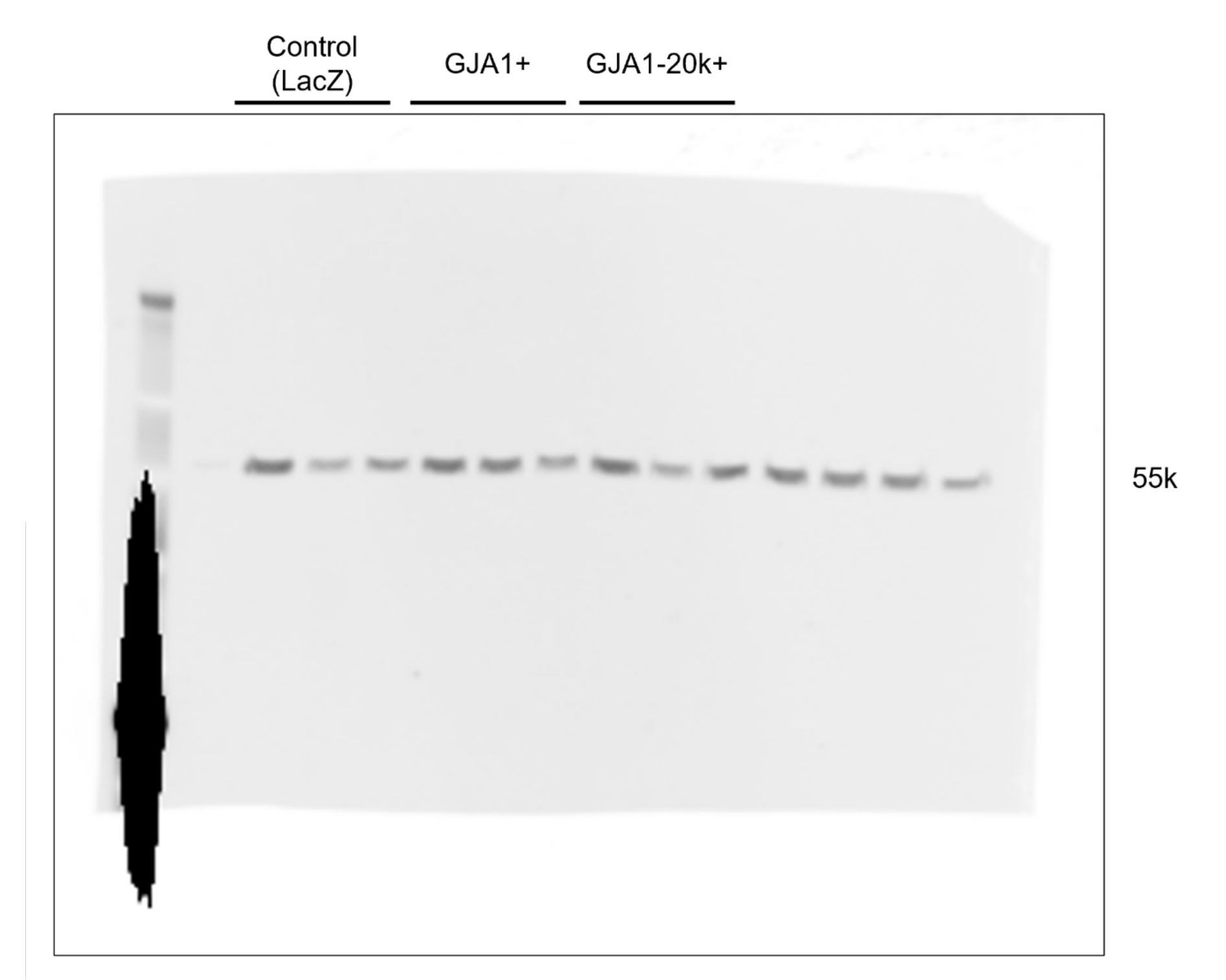
Western blot for LacZ, GJA1+, and GJA1-20k+ lentiviral transduced MSCs: α-tubulin quantification.

**Figure 5.**
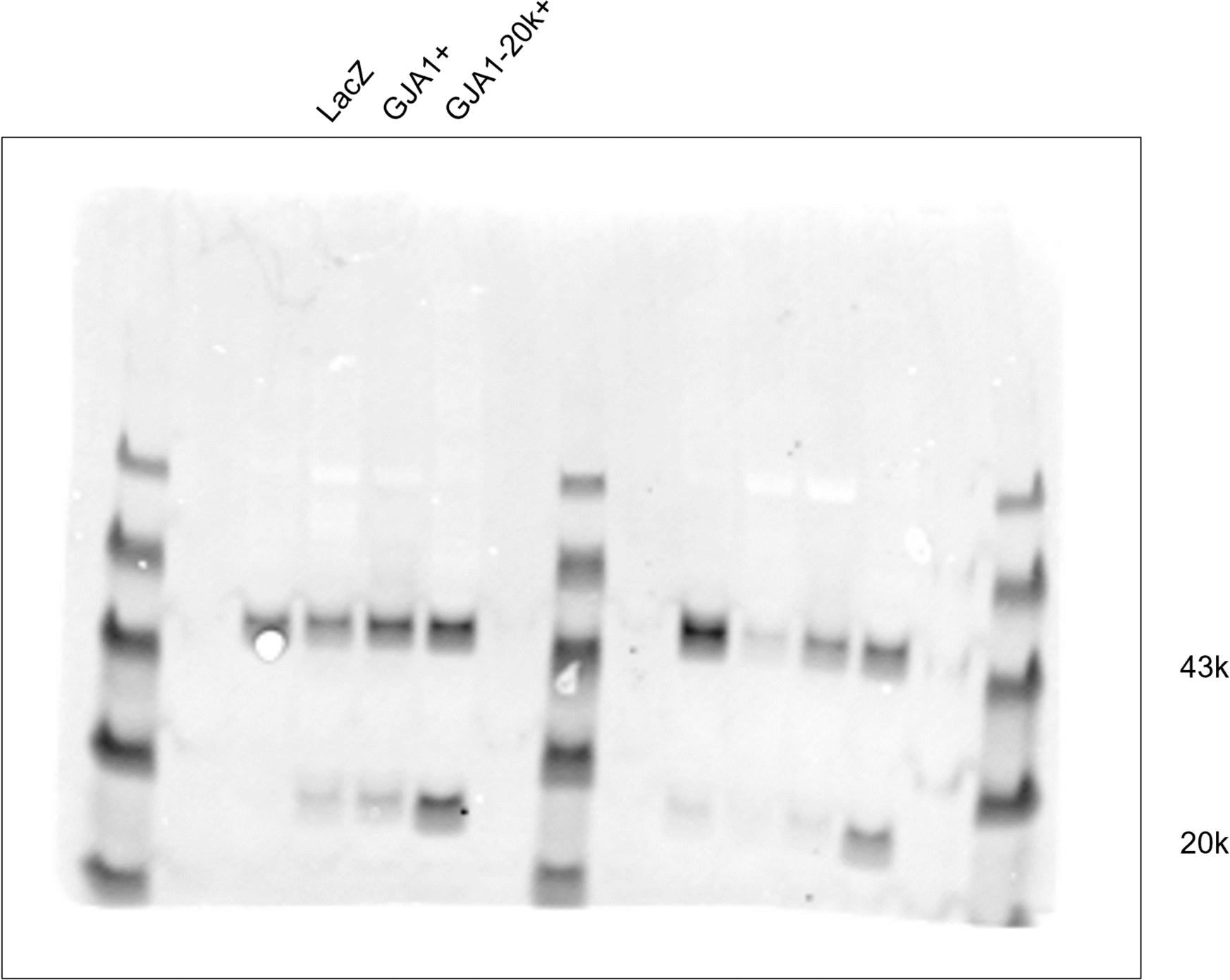
Western blot for LacZ, GJA1+, and GJA1-20k+ lentiviral transduced MSCs: Cx43 single sample representation.

**Figure 6.**
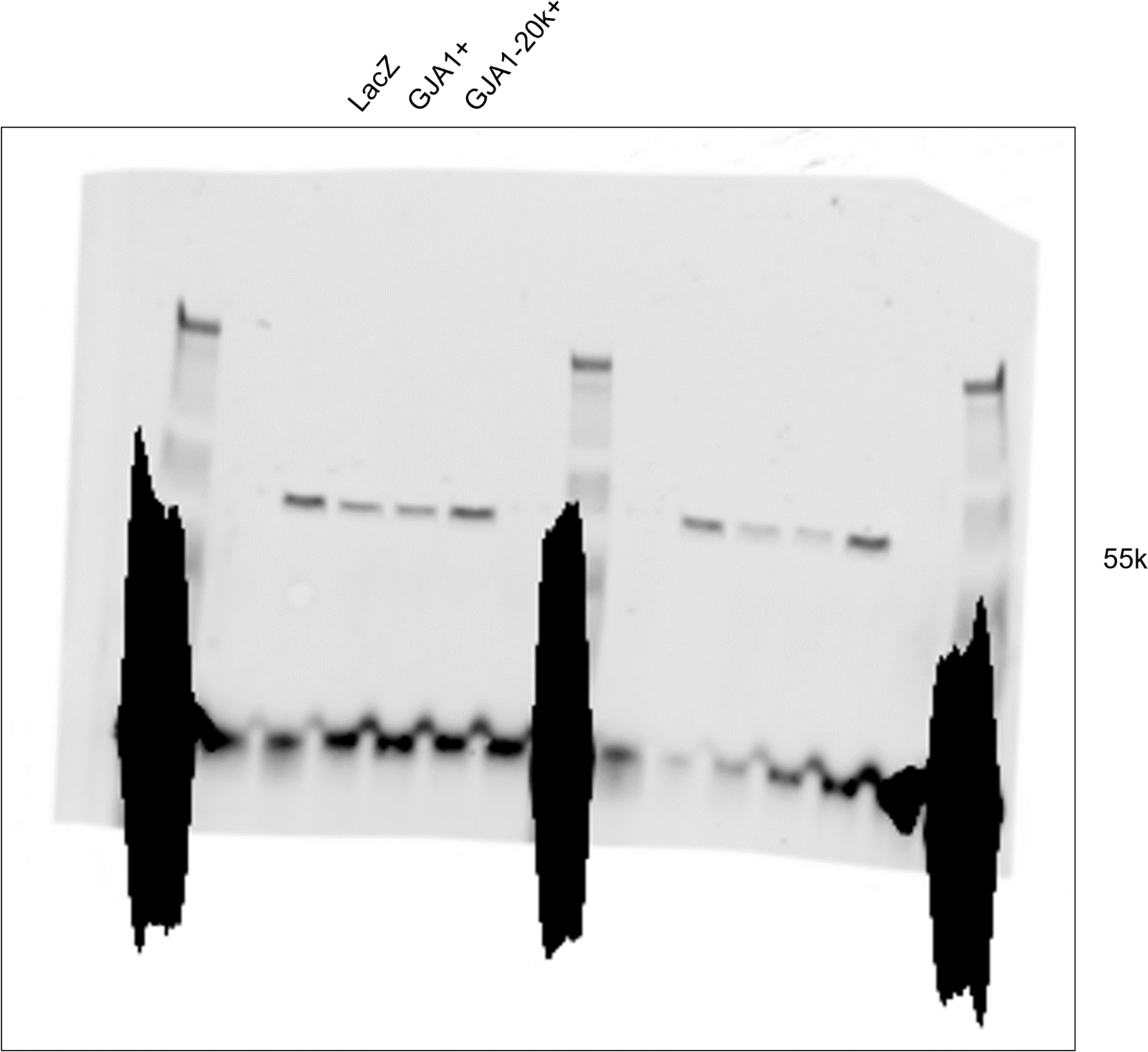
Western blot for LacZ, GJA1+, and GJA1-20k+ lentiviral transduced MSCs: α-tubulin single sample representation.

**Figure 7.**
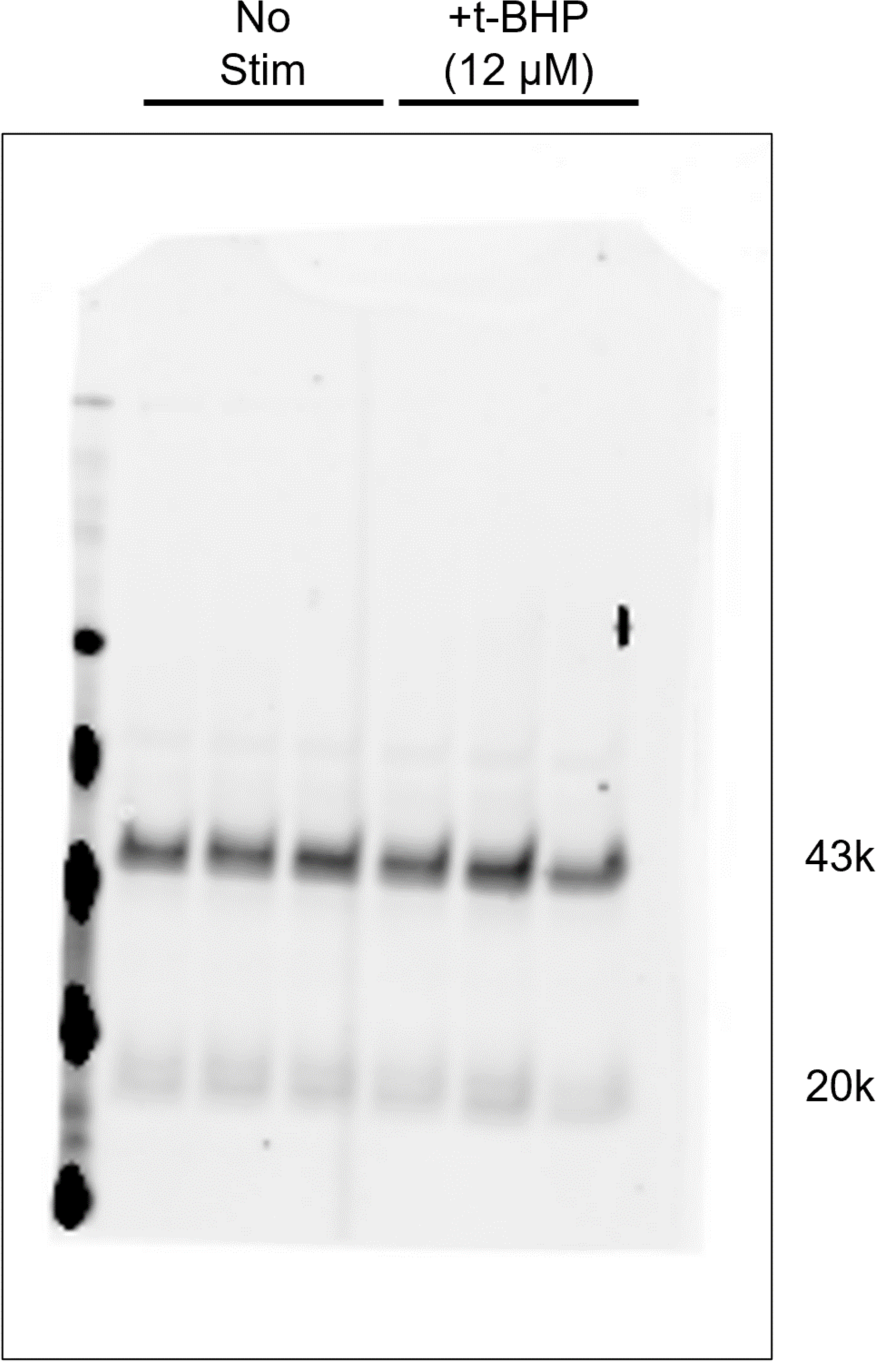
Western blot for chondrocytes +/-t-BHP stimulation: Cx43 quantification.

**Figure 8.**
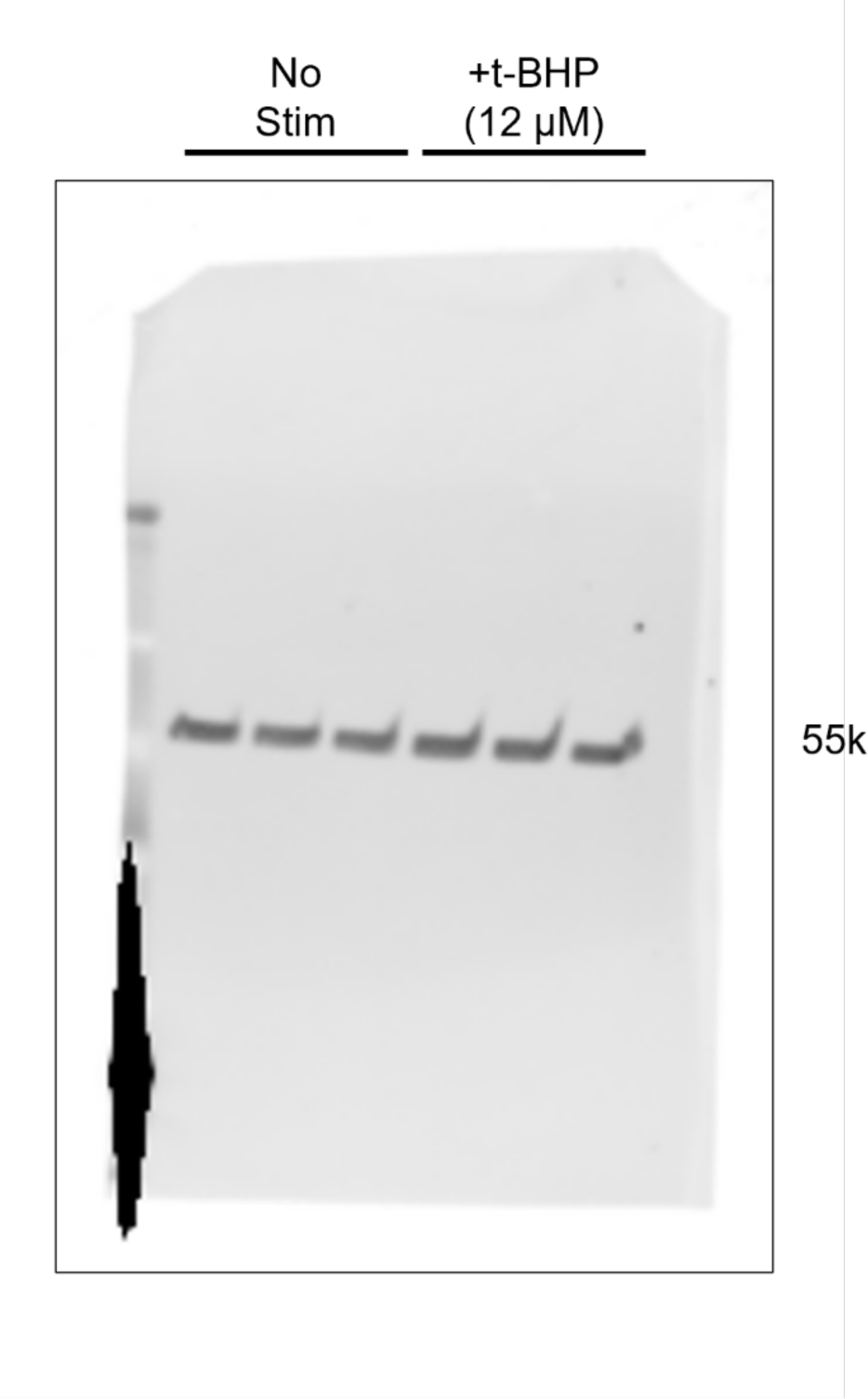
Western blot for chondrocytes +/-t-BHP stimulation: α-tubulin quantification.

## References

1. GBD 2017 Disease and Injury Incidence and Prevalence Collaborators. Global, regional, and national incidence, prevalence, and years lived with disability for 354 diseases and injuries for 195 countries and territories, 1990–2017: a systematic analysis for the Global Burden of Disease Study 2017. Lancet [Internet]. 2018 Nov 10 [cited 2021 Sep 15];392(10159):1789. Available from: /pmc/articles/PMC6227754/

2. DS C, CJ V. Pharmaceutical therapy for osteoarthritis. PM R. 2012 May;4(5 Suppl).

3. B G, FP T, MI H, RP G, MG C, KB F. Chondroprotection and the prevention of osteoarthritis progression of the knee: a systematic review of treatment agents. Am J Sports Med. 2015 Mar 17;43(3):734–44.

4. Delco ML, Goodale M, Talts JF, Pownder SL, Koff MF, Miller AD, et al. Integrin α10β1-Selected Mesenchymal Stem Cells Mitigate the Progression of Osteoarthritis in an Equine Talar Impact Model. American Journal of Sports Medicine. 2020 Mar 1;48(3):612–23.

5. Zhang S, Teo KYW, Chuah SJ, Lai RC, Lim SK, Toh WS. MSC exosomes alleviate temporomandibular joint osteoarthritis by attenuating inflammation and restoring matrix homeostasis. Biomaterials. 2019 Apr 1;200:35–47.

6. Jin Y, Xu M, Zhu H, Dong C, Ji J, Liu Y, et al. Therapeutic effects of bone marrow mesenchymal stem cells-derived exosomes on osteoarthritis. J Cell Mol Med [Internet]. 2021 Oct 1 [cited 2023 Dec 18];25(19):9281. Available from: /pmc/articles/PMC8500984/

7. clinicaltrial.gov. 2024.

8. Matas J, Orrego M, Amenabar D, Infante C, Tapia-Limonchi R, Cadiz MI, et al. Umbilical Cord-Derived Mesenchymal Stromal Cells (MSCs) for Knee Osteoarthritis: Repeated MSC Dosing Is Superior to a Single MSC Dose and to Hyaluronic Acid in a Controlled Randomized Phase I/II Trial. Stem Cells Transl Med [Internet]. 2019 Mar 1 [cited 2023 Oct 16];8(3):215–24. Available from: https://pubmed.ncbi.nlm.nih.gov/30592390/

9. Chahal J, Gómez-Aristizábal A, Shestopaloff K, Bhatt S, Chaboureau A, Fazio A, et al. Bone Marrow Mesenchymal Stromal Cell Treatment in Patients with Osteoarthritis Results in Overall Improvement in Pain and Symptoms and Reduces Synovial Inflammation. Stem Cells Transl Med [Internet]. 2019 Aug 1 [cited 2023 Oct 16];8(8):746–57. Available from: https://pubmed.ncbi.nlm.nih.gov/30964245/

10. Lotfy A, AboQuella NM, Wang H. Mesenchymal stromal/stem cell (MSC)-derived exosomes in clinical trials. Stem Cell Res Ther [Internet]. 2023 [cited 2024 Jan 7];14:66. Available from: http://creativecommons.org/licenses/by/4.0/.TheCreativeCommonsPublicDomainDedicationwai ver

11. Song N, Scholtemeijer M, Shah K. Mesenchymal Stem Cell Immunomodulation: Mechanisms and Therapeutic potential HHS Public Access. Trends Pharmacol Sci. 2020;41(9):653–64.

12. Liesveld JL, Sharma N, Aljitawi OS. Stem cell homing: From physiology to therapeutics. 2020 [cited 2024 Jan 7]; Available from: https://academic.oup.com/stmcls/article/38/10/1241/6430514

13. MN I, SR D, MT E, M W, L S, K W, et al. Mitochondrial transfer from bone-marrow-derived stromal cells to pulmonary alveoli protects against acute lung injury. Nat Med [Internet]. 2012 May [cited 2021 Sep 15];18(5):759–65. Available from: https://pubmed.ncbi.nlm.nih.gov/22504485/

14. Wang J, Li H, Yao Y, Zhao T, Chen YY, Shen YL, et al. Stem cell-derived mitochondria transplantation: A novel strategy and the challenges for the treatment of tissue injury. Stem Cell Res Ther. 2018 Apr 13;9(1).

15. Morrison TJ, Jackson M V., Cunningham EK, Kissenpfennig A, McAuley DF, O’Kane CM, et al. Mesenchymal stromal cells modulate macrophages in clinically relevant lung injury models by extracellular vesicle mitochondrial transfer. Am J Respir Crit Care Med [Internet]. 2017 Nov 15 [cited 2023 Oct 15];196(10):1275–86. Available from: www.atsjournals.org

16. Jackson M V, Morrison TJ, Doherty DF, Mcauley DF, Matthay MA, Kissenpfennig A, et al. Mitochondrial Transfer via Tunneling Nanotubes is an Important Mechanism by Which Mesenchymal Stem Cells Enhance Macrophage Phagocytosis in the In Vitro and In Vivo Models of ARDS SIGNIFICANCE STATEMENT. Stem Cells [Internet]. 2016 [cited 2023 Oct 15];34:2210–23. Available from: www.listlabs.com

17. Acquistapace A, Bru T, Ois Lesault P franç, Figeac F, Lie Coudert AE, Coz O LE, et al. Human Mesenchymal Stem Cells Reprogram Adult Cardiomyocytes Toward a Progenitor-Like State Through Partial Cell Fusion and Mitochondria Transfer. Stem Cells [Internet]. 2011 [cited 2023 Oct 15];29:812–24. Available from: www.abcam.

18. Babenko VA, Silachev DN, Zorova LD, Pevzner IB, Khutornenko AA, Plotnikov EY, et al. Improving the Post-Stroke Therapeutic Potency of Mesenchymal Multipotent Stromal Cells by Cocultivation With Cortical Neurons: The Role of Crosstalk Between Cells. Stem Cells Transl Med [Internet]. 2015 Sep 1 [cited 2023 Oct 15];4(9):1011–20. Available from: 10.5966/sctm.2015-0010

19. Babenko VA, Silachev DN, Popkov VA, Zorova LD, Pevzner IB, Plotnikov EY, et al. Miro1 Enhances Mitochondria Transfer from Multipotent Mesenchymal Stem Cells (MMSC) to Neural Cells and Improves the Efficacy of Cell Recovery. Molecules [Internet]. 2018 [cited 2023 Oct 16];23(3). Available from: https://pubmed.ncbi.nlm.nih.gov/29562677/

20. Shen J, Zhang JH, Xiao H, Wu JM, He KM, Lv ZZ, et al. Mitochondria are transported along microtubules in membrane nanotubes to rescue distressed cardiomyocytes from apoptosis. Cell Death Dis [Internet]. 2018 Feb 1 [cited 2023 Oct 16];9(2). Available from: https://pubmed.ncbi.nlm.nih.gov/29362447/

21. Liu D, Gao Y, Liu J, Huang Y, Yin J, Feng Y, et al. Intercellular mitochondrial transfer as a means of tissue revitalization. Signal Transduct Target Ther. 2021 Dec 1;6(1).

22. Zhang Y, Yu Z, Jiang D, Liang X, Liao S, Zhang Z, et al. iPSC-MSCs with High Intrinsic MIRO1 and Sensitivity to TNF-α Yield Efficacious Mitochondrial Transfer to Rescue Anthracycline-Induced Cardiomyopathy. Stem Cell Reports [Internet]. 2016 Oct 11 [cited 2023 Oct 16];7(4):749–63. Available from: https://pubmed.ncbi.nlm.nih.gov/27641650/

23. Konari N, Nagaishi K, Kikuchi S, Fujimiya M. Mitochondria transfer from mesenchymal stem cells structurally and functionally repairs renal proximal tubular epithelial cells in diabetic nephropathy in vivo. Sci Rep. 2019 Dec 1;9(1).

24. Wei B, Ji M, Lin Y, Wang S, Liu Y, Geng R, et al. Mitochondrial transfer from bone mesenchymal stem cells protects against tendinopathy both in vitro and in vivo. Stem Cell Res Ther. 2023 Dec 1;14(1).

25. Fahey M, Bennett M, Thomas M, Montney K, Vivancos-Koopman I, Pugliese B, et al. Mesenchymal stromal cells donate mitochondria to articular chondrocytes exposed to mitochondrial, environmental, and mechanical stress. Sci Rep [Internet]. 2022 Dec 1 [cited 2023 Oct 15];12(1). Available from: https://pubmed.ncbi.nlm.nih.gov/36513773/

26. Korpershoek J V., Rikkers M, Wallis FSA, Dijkstra K, te Raa M, de Knijff P, et al. Mitochondrial Transport from Mesenchymal Stromal Cells to Chondrocytes Increases DNA Content and Proteoglycan Deposition In Vitro in 3D Cultures. Cartilage. 2022 Dec 1;13(4):133–47.

27. Wang R, Maimaitijuma T, Ma YY, Jiao Y, Cao YP. Mitochondrial transfer from bone-marrow-derived mesenchymal stromal cells to chondrocytes protects against cartilage degenerative mitochondrial dysfunction in rats chondrocytes. Chin Med J (Engl) [Internet]. 2020 Jan 20 [cited 2022 Apr 3];134(2):212–8. Available from: https://pubmed.ncbi.nlm.nih.gov/32858593/

28. Delco ML, Bonnevie ED, Bonassar LJ, Fortier LA. Mitochondrial dysfunction is an acute response of articular chondrocytes to mechanical injury. Journal of Orthopaedic Research [Internet]. 2018 Feb 1 [cited 2020 Feb 26];36(2):739–50. Available from: http://doi.wiley.com/10.1002/jor.23651

29. Delco ML, Bonnevie ED, Szeto HS, Bonassar LJ, Fortier LA. Mitoprotective therapy preserves chondrocyte viability and prevents cartilage degeneration in an ex vivo model of posttraumatic osteoarthritis. Journal of Orthopaedic Research. 2018 Aug 1;36(8):2147–56.

30. LR B, LA F, LJ B, HH S, I C, ML D. Mitoprotective therapy prevents rapid, strain-dependent mitochondrial dysfunction after articular cartilage injury. J Orthop Res [Internet]. 2020 Jun 1 [cited 2021 Sep 15];38(6):1257–67. Available from: https://pubmed.ncbi.nlm.nih.gov/31840828/

31. Y Y, XL F, D J, Y Z, X L, ZB X, et al. Connexin 43-Mediated Mitochondrial Transfer of iPSC-MSCs Alleviates Asthma Inflammation. Stem Cell Reports [Internet]. 2018 Nov 13 [cited 2021 Sep 15];11(5):1120–35. Available from: https://pubmed.ncbi.nlm.nih.gov/30344008/

32. Osswald M, Jung E, Sahm F, Solecki G, Venkataramani V, Blaes J, et al. Brain tumour cells interconnect to a functional and resistant network. Nature [Internet]. 2015 Dec 3 [cited 2023 Oct 16];528(7580):93–8. Available from: https://pubmed.ncbi.nlm.nih.gov/26536111/

33. Norris RP, Rachael Norris CP. Transfer of mitochondria and endosomes between cells by gap junction internalization. Traffic [Internet]. 2021 [cited 2023 Mar 14];22:174–9. Available from: https://onlinelibrary.wiley.com/doi/10.1111/tra.12786

34. Golan K, Singh AK, Kollet O, Bertagna M, Althoff MJ, Khatib-Massalha E, et al. Bone marrow regeneration requires mitochondrial transfer from donor Cx43-expressing hematopoietic progenitors to stroma. Blood [Internet]. 2020 Dec 3 [cited 2023 Oct 16];136(23):2607–19. Available from: https://pubmed.ncbi.nlm.nih.gov/32929449/

35. Yang J, Liu L, Oda Y, Wada K, Ago M, Matsuda S, et al. Extracellular Vesicles and Cx43-Gap Junction Channels Are the Main Routes for Mitochondrial Transfer from Ultra-Purified Mesenchymal Stem Cells, RECs. Int J Mol Sci [Internet]. 2023 Jun 1 [cited 2023 Oct 16];24(12). Available from: https://pubmed.ncbi.nlm.nih.gov/37373439/

36. Smyth JW, Shaw RM. Autoregulation of connexin43 gap junction formation by internally translated isoforms. Cell Rep [Internet]. 2013 [cited 2022 Jan 10];5(3):611–8. Available from: https://pubmed.ncbi.nlm.nih.gov/24210816/

37. Y F, SS Z, S X, WA B, R B, I E, et al. Cx43 Isoform GJA1-20k Promotes Microtubule Dependent Mitochondrial Transport. Front Physiol [Internet]. 2017 Nov 7 [cited 2021 Sep 15];8(NOV). Available from: https://pubmed.ncbi.nlm.nih.gov/29163229/

38. Basheer WA, Xiao S, Epifantseva I, Fu Y, Kleber AG, Hong TT, et al. GJA1-20k Arranges Actin to Guide Cx43 Delivery to Cardiac Intercalated Discs. Circ Res. 2017 Oct 13;121(9):1069–80.

39. Ren D, Zheng P, Zou S, Gong Y, Wang Y, Duan J, et al. GJA1-20K Enhances Mitochondria Transfer from Astrocytes to Neurons via Cx43-TnTs After Traumatic Brain Injury. Cell Mol Neurobiol [Internet]. 2021; Available from: 10.1007/s10571-021-01070-x

40. James CC, Zeitz MJ, Calhoun PJ, Lamouille S, Smyth JW. Altered translation initiation of Gja1 limits gap junction formation during epithelial-mesenchymal transition. Mol Biol Cell [Internet]. 2018 Apr 1 [cited 2022 Sep 29];29(7):797–808. Available from: https://pubmed.ncbi.nlm.nih.gov/29467255/

41. R GF, P CF, MB G, PR B, MD M, FJ B. Biochemical evidence for gap junctions and Cx43 expression in immortalized human chondrocyte cell line: a potential model in the study of cell communication in human chondrocytes. Osteoarthritis Cartilage [Internet]. 2014 [cited 2021 Sep 22];22(4):586–90. Available from: https://pubmed.ncbi.nlm.nih.gov/24530659/

42. Yang J, Liu L, Oda Y, Wada K, Ago M, Matsuda S, et al. Highly-purified rapidly expanding clones, RECs, are superior for functional-mitochondrial transfer. Stem Cell Res Ther [Internet]. 2023 Dec 1 [cited 2023 Oct 19];14(1):1–22. Available from: https://stemcellres.biomedcentral.com/articles/10.1186/s13287-023-03274-y

43. Shimura D, Nuebel E, Baum R, Valdez SE, Xiao S, Warren JS, et al. Protective mitochondrial fission induced by stress-responsive protein GJA1-20k. Elife. 2021;

44. Basheer WA, Fu Y, Shimura D, Xiao S, Agvanian S, Hernandez DM, et al. Stress response protein GJA1-20k promotes mitochondrial biogenesis, metabolic quiescence, and cardioprotection against ischemia/reperfusion injury. JCI Insight. 2018 Oct 18;3(20).

45. Thomas MA, Fahey MJ, Pugliese BR, Irwin RM, Antonyak MA, Delco ML. Human mesenchymal stromal cells release functional mitochondria in extracellular vesicles. Front Bioeng Biotechnol [Internet]. 2022 Aug 19 [cited 2022 Dec 12];10:870193. Available from: http://www.ncbi.nlm.nih.gov/pubmed/36082164

46. Fu YL, Tao L, Peng FH, Zheng NZ, Lin Q, Cai SY, et al. GJA1-20k attenuates Ang II-induced pathological cardiac hypertrophy by regulating gap junction formation and mitochondrial function. Acta Pharmacol Sin [Internet]. 2020; Available from: 10.1038/s41401-020-0459-6

47. Md M, P Cf, R Gf, O M de I, Hz W, V V, et al. Human articular chondrocytes express multiple gap junction proteins: differential expression of connexins in normal and osteoarthritic cartilage. Am J Pathol [Internet]. 2013 Apr [cited 2021 Sep 15];182(4):1337–46. Available from: https://pubmed.ncbi.nlm.nih.gov/23416160/

48. Zeitz MJ, Calhoun PJ, James CC, Taetzsch T, George KK, Robel S, et al. Dynamic UTR Usage Regulates Alternative Translation to Modulate Gap Junction Formation during Stress and Aging. Cell Rep. 2019 May 28;27(9):2737–2747.e5.

49. Salat-Canela C, Sesé M, Peula C, Ramón Y Cajal S, Aasen T. Internal translation of the connexin 43 transcript. Cell Commun Signal [Internet]. 2014 May 8 [cited 2023 Dec 20];12(1):31. Available from: /pmc/articles/PMC4108066/

50. Andrew Sinclair K, Terase Yerkovich S, Mark-Anthony Hopkins P, Charles Chambers D. Characterization of intercellular communication and mitochondrial donation by mesenchymal stromal cells derived from the human lung. 2016;

